# A gene-based model of fitness and its implications for genetic variation: Genetic and inbreeding loads

**DOI:** 10.1101/2025.02.19.639162

**Authors:** Parul Johri, Brian Charlesworth

## Abstract

In the companion paper to this, we examined the consequences for patterns of linkage disequilibrium of the “gene” model of fitness, which postulates that the effects of recessive or partially recessive deleterious mutations located at different sites within a gene fail to complement each other. Here, we examine the consequences of the gene model for the genetic and inbreeding loads, using both analytical and simulation methods, and contrast it with the frequently used “sites” model that allows allelic complementation. We show that the gene model results in a slightly lower genetic load, but a much smaller inbreeding load, than the sites model, implying that standard predictions of mutational contributions to inbreeding depression may be overestimates. Synergistic epistasis between pairs of mutations was also modeled, and shown to considerably reduce the inbreeding load for both the gene and sites models. The theoretical results are discussed in relation to data on inbreeding load in *Drosophila melanogaster*. The widespread assumption that inbreeding depression is largely due to deleterious mutations should be re-examined in the light of our findings.

## Introduction

A large volume of recent work on the properties of diploid populations subject to the input of deleterious mutations has been stimulated by concerns about their implications for the survival of small, endangered populations, and by the availability of population genetic data on non-model organisms that shed light on the prevalence of deleterious mutations in natural populations (e.g., Teixeira and Huber 2021; Kardos et al. 2021; Bertorelli et al. 2022; Peréz-Pereira et al. 2002; Kyriazis, Robinson and Lohmueller 2023; Robinson et al. 2023). There are, however, sharp disagreements about the interpretation of the data (Kyriazis et al., 2021, 2023; García-Dorado and Hedrick 2023), emphasizing the need for a secure theoretical basis for data analyses.

In the companion paper to this (Johri and Charlesworth 2025), we analysed the consequences for patterns of linkage disequilibrium (LD) of the differences between two models of the fitness effects of deleterious mutations located within the same gene, the “gene” and “sites” models. The gene model assumes that two heterozygous mutations in *trans* have a larger effect on fitness than two mutations in *cis*, due to a lack of complementation, whereas the sites model assumes that there is no such difference. Many theoretical predictions about the properties of deleterious mutations in populations, and their effects on features such as population mean fitness, genetic variance in fitness and inbreeding depression have been based on the sites model, together with the assumption of multiplicative fitnesses when the effects of mutations at different sites are combined (for a recent review, see Kyriazis, Robinson and Lohmueller 2023).

But the difference between the gene and sites models can affect the effective level of recessivity experienced by deleterious mutations, as can be seen from the following, highly simplified, example. Consider a single coding sequence that is segregating for mutations at two different nucleotide sites (1 and 2) with the same selection coefficient *s* against homozygotes at each site. Assume that the double mutant haplotypes are sufficiently rare that they can be ignored. Let the frequency of mutant variants at site *i* be *q_i_*. Then haplotypes carrying a mutation at site 1 will meet a haplotype that is wild-type at site 1 and mutant at site 2 with an approximate frequency of *p*_1_*q*_2_; the fitness of this genotype is 1 – *s* if there is no allelic complementation (*i.e*, under the gene model), but the sites model would assign it a fitness of 1 – 2*hs* (assuming additive effects across sites), where *h* is the dominance coefficient. It will meet haplotypes that are mutant at site 1 but wild-type at site 2 with a frequency of approximately *q*_1_*p*_2_. Both models would assign fitness 1 – *s* to this genotype. A similar argument applies to a haplotype carrying a mutation at site 2, with the appropriate switch between subscripts 1 and 2.

All of the double mutant genotypes are classed as homozygotes under the gene model, which is biologically realistic for the majority of genes, for which sites with allelic complementation are relatively rare (Fincham et al. 1979, pp.348-353; Hawley and Gilliland 2006). The mean difference between the gene and sites models in the fitness reduction assigned to haplotypes that each carry a mutation at one or other site is thus:

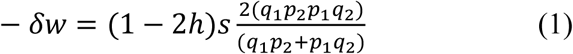

If the frequencies of the deleterious mutations at the two sites are similar, so that the subscripts can be dropped, – *δw* ≈ 2(1 − 2*h*)*spq*, which is positive if *h* < < 0.5. If *h* > 0 and the population is at deterministic mutation-selection balance, *q* ≈ *u*/(*hs*), (Haldane 1927), and − *δw* ≈ 2(1 − 2*h*)*u*/*h*. For completely recessive deleterious mutations at equilibrium, *q* ≈ 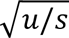 and −*δw* ≈ 2(1 − 2*h*)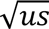. This implies that the sites model can underestimate the true homozygous fitness effects of deleterious mutations, although the effect is small for a single pair of rare deleterious variants. It follows that deleterious mutations are more likely to segregate at higher allele frequencies under the sites model than the gene model, leading to a lower mean fitness of the population, a higher genetic load, and a larger effect of inbreeding on fitness. In addition to this effect, our previous paper showed that the gene model can generate positive LD among deleterious variants under circumstances where the sites model causes zero or slightly negative LD (Johri and Charlesworth 2025). Since positive LD enhances the efficacy of selection (e.g., Barton 1995; Roze 2021), this effect would also help to reduce the frequency of deleterious mutations expected under the gene model compared with the sites model.

The purpose of the present paper is to examine the consequences of the differences between the gene and sites models for the load statistics, using deterministic and stochastic simulation models of a single coding sequence subject to deleterious mutations, which were described in Johri and Charlesworth (2025). We show that the difference between the two models has especially significant consequences for the level of inbreeding depression caused by deleterious mutations, suggesting that predictions based on the sites model could greatly overestimate the expected level of inbreeding depression contributed by deleterious mutations. We also show that a model of pairwise synergistic epistasis between mutations within the same gene results in even lower levels of inbreeding depression than in its absence. We also note that the sites model is used in current methods for estimating the distribution of fitness effects of deleterious mutations, and that this will lead to overestimation of the mean strength of selection against such mutations if the gene model is more appropriate.

## Methods

### Fitness models and simulation methods

The fitness models and simulation results used here are described in the Methods section of Johri and Charlesworth (2025). Recursion relations for haplotype frequencies for the two-site deterministic model are described in the Appendix of that paper; use of these expressions allows the numerical values of the equilibrium haplotype frequencies to be determined for a given set of mutation and fitness parameters. In addition, the multi-site approximation for the gene model without epistasis with the fixed selection and dominance coefficients at all sites, described in the penultimate section of the Appendix to Johri and Charlesworth (2025), provides expressions for the load statistics that can be compared with the simulations when selection is strong relative to genetic drift. Standard formulae for mutation-selection balance can be used for the corresponding calculations with the sites model (see section 2 of the Appendix to the present paper).

### Estimating the load statistics from the simulations

The state of a population with *N* diploid individuals (where *N* was usually equal to 1000) with respect to the load statistics was summarized using four quantities – the genetic load (*L*), the inbreeding load (*B*), the variance in fitness between individuals (*V*), and the mean frequency of mutant alleles per site (*q̅*). The genetic load for each simulation replicate was calculated as *L* = 1 − *w̅*, where *w̅* is the mean fitness of the population for all individuals in the simulated population, relative to a value of 1 for individuals homozygous for the wild-type allele at all sites; *w̅* was calculated using all individuals sampled after burn-in. The variance *V* was calculated in a similar way. Allele frequencies (*q*) were estimated from a sample of 100 genomes and included monomorphic sites.

To calculate *B*, which is defined here as the mean difference in fitness between completely outbred and completely inbred individuals, 100 genomes were sampled randomly from the population, from which 100 diploid inbred individuals were artificially created (*i.e*, each haploid genome was duplicated from a single diploid individual). The inbreeding load was calculated as *B* = *w̅* − *w̅*_*I*_, where *w̅*_*I*_ is the mean fitness of the subsampled inbred population. Because *L* and *B* are both small, this expression for *B* differs little from the usual expression for inbreeding depression, 1 − *w̅*_*I*_/*w̅*. Under the assumption of small multiplicative fitness effects across sites, it is equivalent to the negative of the regression coefficient of the natural logarithm of fitness on inbreeding coefficient (Morton et al. 1956).

Each simulation replicate yields a single data point for each variable, and the distributions, means and standard errors shown in the figures and tables are derived from 1000 replicate simulations.

### Data availability

The scripts used to determine the properties of two-site equilibrium populations, perform all the simulations, and calculate population genetic statistics are provided at https://github.com/paruljohri/Gene_vs_sites_model/tree/main.

## Results

### Deterministic results for two selected sites

To aid the reader, Table 1 of Johri and Charlesworth (2025) is reproduced here. The fact that the two sites have identical fitness effects means that the four haplotype frequencies, *x*_1_ (+ +), *x*_2_ (– +), *x*_3_ (+ –) and *x*_4_ (– –) (where + and – indicates wild-type and mutant alleles, respectively) can be replaced with *x* = *x*_2_ = *x*_3_, *y* = *x*_4_, and *z* =1 – 2*x* – *y*. The frequency of the – allele at a site is *q* = *x* + *y*. The load statistics *L*, *B,* and *V* (the genetic load, the inbreeding load, and the genetic variance, respectively) can easily be determined once the equilibrium haplotype frequencies are known, using the formulae presented in the Appendix, Equations (A1) – A(8). As in Johri and Charlesworth (2025), we assume no recombination between the two sites, which should provide a good approximation to what is expected for a single gene.

**Table 1.** Models of the fitness effects of two sites with a fixed selection coefficient.

|  | ++ | + - | - + | -- |
| --- | --- | --- | --- | --- |
| ++ | 1 | $1 - hs$ | $1 - hs$ | $1 - 2hs - e_1$ |
| - + | $1 - hs$ | $1 - 2hs - e_1$ | $1 - s$ | $1 - s - hs - e_2$ |
| + - | $1 - hs$ | $1 - s$ | $1 - 2hs - e_1$ | $1 - s - hs - e_2$ |
| -- | $1 - 2hs - e_1$ | $1 - s - hs - e_2$ | $1 - s - hs - e_2$ | $1 - 2s - e_3$ |

$$e_1 = 2\epsilon hs; e_2 = \epsilon(1 + h)s; e_3 = 2\epsilon s.$$ The “gene” model
|  | ++ | + - | - + | -- |
| --- | --- | --- | --- | --- |
| ++ | 1 | $1 - hs$ | $1 - hs$ | $1 - 2hs - e_1$ |
| - + | $1 - hs$ | $1 - s$ | $1 - s$ | $1 - 1.5s - e_2$ |
| + - | $1 - hs$ | $1 - s$ | $1 - s$ | $1 - 1.5s - e_2$ |
| -- | $1 - 2hs - e_1$ | $1 - 1.5s - e_2$ | $1 - 1.5s - e_2$ | $1 - 2s - e_3$ |

Note that the parameter $k$ used in the gene model in Table 1 of Johri and Charlesworth (2025) is here set to $\frac{1}{2}$ .

Table 2 shows the results for the case of *u* = 0.0001, *s* = 0.01, which was previously used for the LD statistics presented in Johri and Charlesworth (2025). As noted there, these are not realistic parameter values, but produce relatively large equilibrium allele frequencies, enhancing the contrast between the two models. The frequency of deleterious mutations (*q_r_* in Table 2) is represented by the ratio of the exact deterministic equilibrium value to the approximate value for a single site subject to mutation and selection, given by Equation (A4) of Johri and Charlesworth (2025). Since the denominator *q_r_* is independent of the strength of epistasis, and is the same for the gene and sites model, differences in *q_r_* between the two models and between different values of the epistasis coefficient *ε* for a given *h* and *s* reflect differences in the absolute deleterious allele frequency, *q*.

**Table 2.**
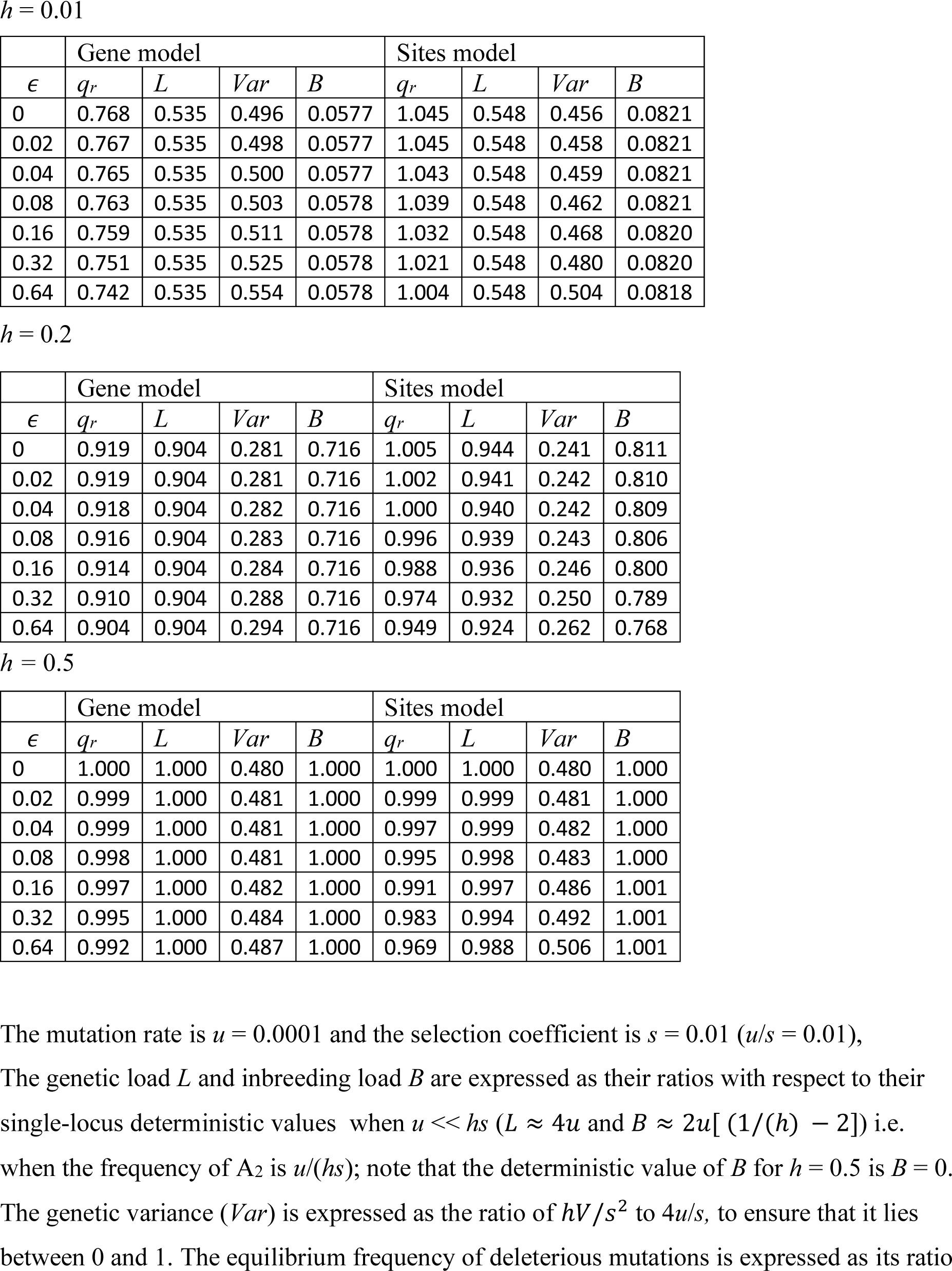
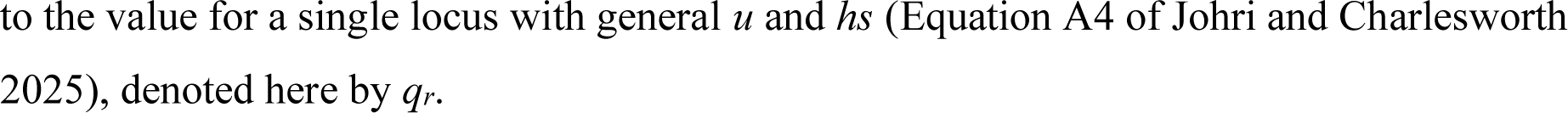
Population genetic statistics for the gene and sites models with different values of the dominance coefficient (*h*) and epistasis coefficient (*ε*)

*L* and *B* are represented by the ratios of their values to those for an additive model of two independent loci, when *h* > 0 and *u* << *hs*. In this case, *L* ≈ 4*u* (see Haldane 1937) and *B* ≈ 2*u*[ (1/(*h*) − 2] (see Morton et al. 1956); if *u* is sufficiently large relative to *hs* or drift is operating, these formulae no longer hold. To avoid large values when *h* is small, the variance is represented by *Var* = *hV* /(4*uhs*), the denominator being the equilibrium value of *V* for the purely additive case with no LD and *u* << *hs*. This means that results for the variances with different *h* values are not directly comparable, but the sites and gene models with the same *h* can be compared.

As before, we focus attention on cases with zero epistasis (*ε* = 0) and synergistic epistasis (*ε* > 0). Over the range of values of *ε* considered here, its magnitude has relatively little effect on *L* and *B,* for both the sites and gene models. Its effect on *q_r_* is more complex. For both models, *q* decreases with *ε*, but the decline is faster for the sites model when *h* is close to ½, with the result that in this case *q* is smaller for the sites model than the gene model when *ε* is large, despite their having the same *q* with no epistasis. For *h* sufficiently smaller than ½, *q* is smaller for the gene model.

The gene model, as might be expected, is associated with lower values of *L* and *B* than the sites model when *h* <½, with a much lower value of *B* than the sites model when h is small. The higher values of *V* under the gene model with small *h* reflect its larger values of LD compared with the sites model, with positive associations (*D* > 0) between deleterious variants when *ε* is sufficiently small, especially with small *h* (Johri and Charlesworth 2025). With *h* = ½, *B* = 0 (so that the ratios in Table 2 are set to one) and *L* is the same for the two models, as expected. *V* increases more quickly with *ε* for the sites model than the gene model, the reverse of the pattern for *q*. This can result in a slightly higher *V* for the sites model with large *ε* and *h*, probably reflecting the larger contributions of terms in *εhs* under the sites model, as can be seen in the fitness matrices in Table 1.

### Simulation results for multiple sites

#### Load statistics with no epistasis

The analytical results described above suggest that, relative to the sites model, the gene model should result in smaller mean frequencies of the deleterious alleles, causing a a lower genetic load and inbreeding load. The multiple site simulations with scaled parameters that are appropriate for *Drosophila* populations, *i.e*, 1000 selected sites, *Nu* = 0.005, and *Nr* = 0.01 (for details, see the *Methods* section of Johri and Charlesworth 2025), show that the equilibrium frequencies of deleterious alleles are much smaller for the gene than the sites model, except when selection is weak relative to drift (*γ* = 2 for the case of a fixed selection coefficient (Table 3) and 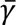 = 2 for the case of a gamma distribution of selection coefficients (Table 4), in which case there is little difference between the two models. There is also a drastic difference between the estimated inbreeding load (*B*) between the two models: *B* is much smaller for the gene than the sites model when *h* < ½, even with weak selection. There is a much smaller effect on the genetic load (*L*), which is noticeable only when mutations are fully recessive.

**Table 3.**
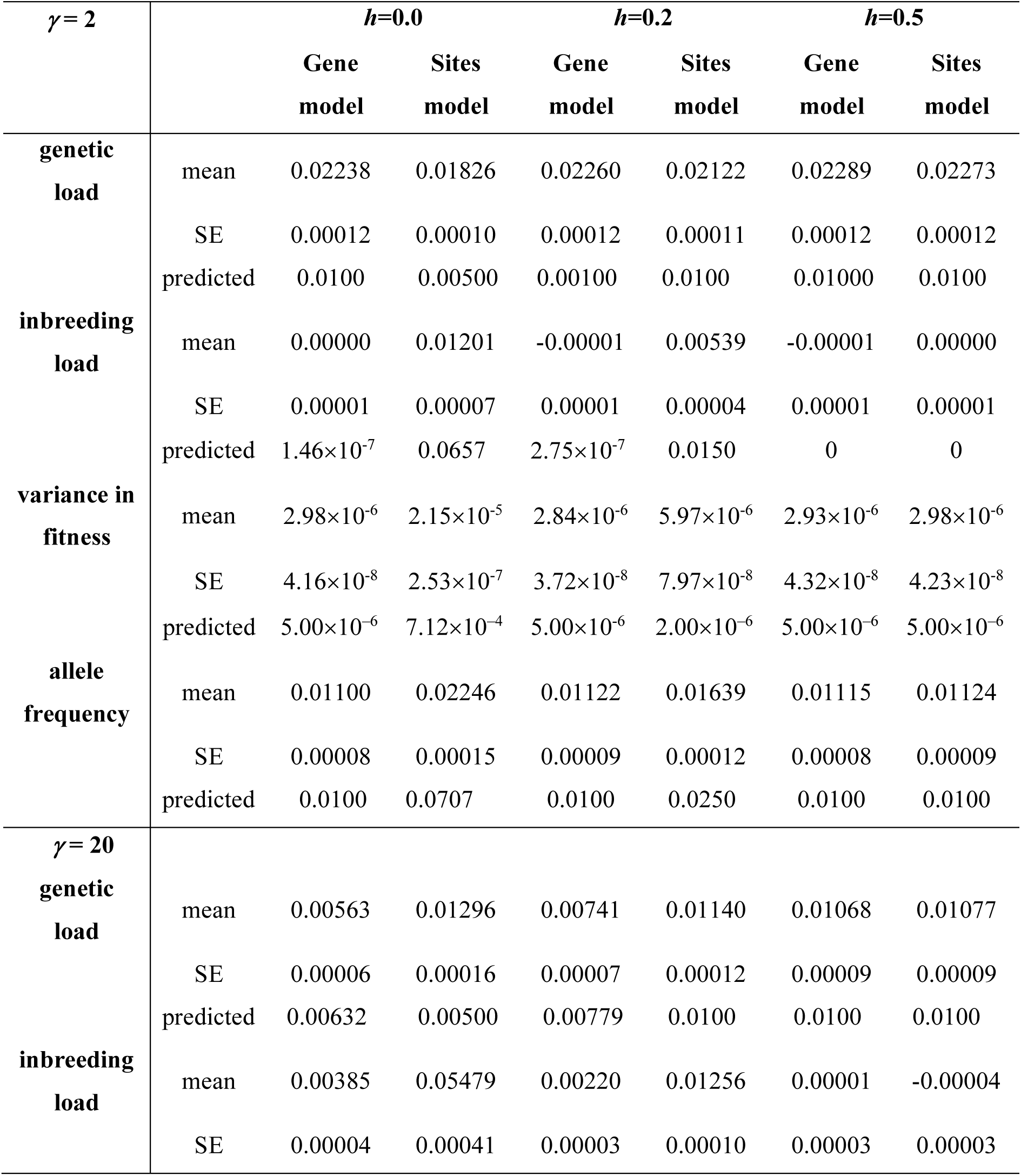
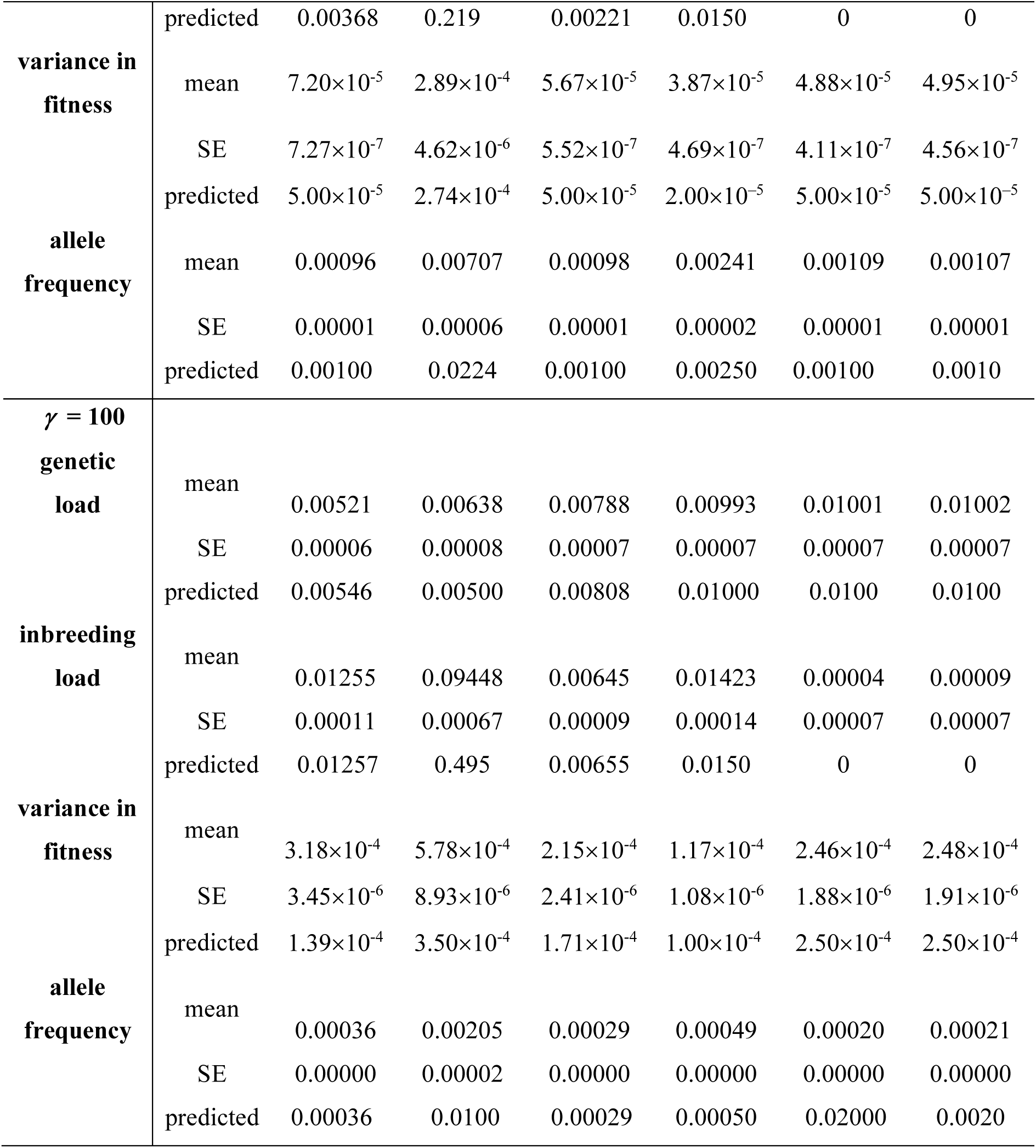
The load statistics for the gene and sites models with three fixed values of *γ* = 2*Ns.* 1000 selected sites were simulated, with *Nu* = 0.005 and *Nr* = 0.01, where *N* is the population size (*N* = 1000), *u* is the mutation rate per site/generation, and *r* is the rate of crossing over between adjacent sites. Details of how to estimate the load statistics are given in the Methods section. The means and standard errors (SEs) for 1000 replicate simulations are shown. The predicted values obtained by the methods described in the text are also shown.

**Table 4.**
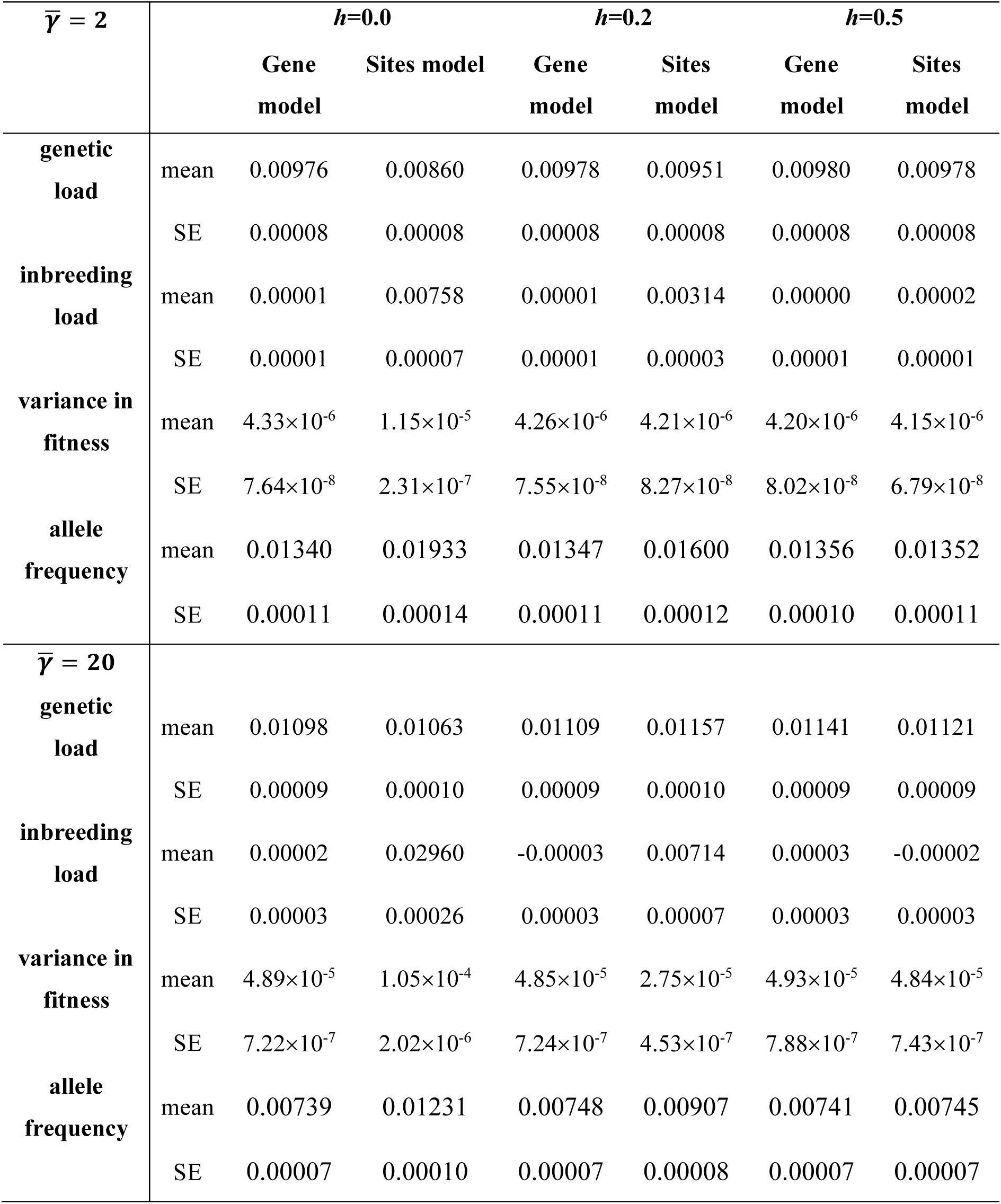
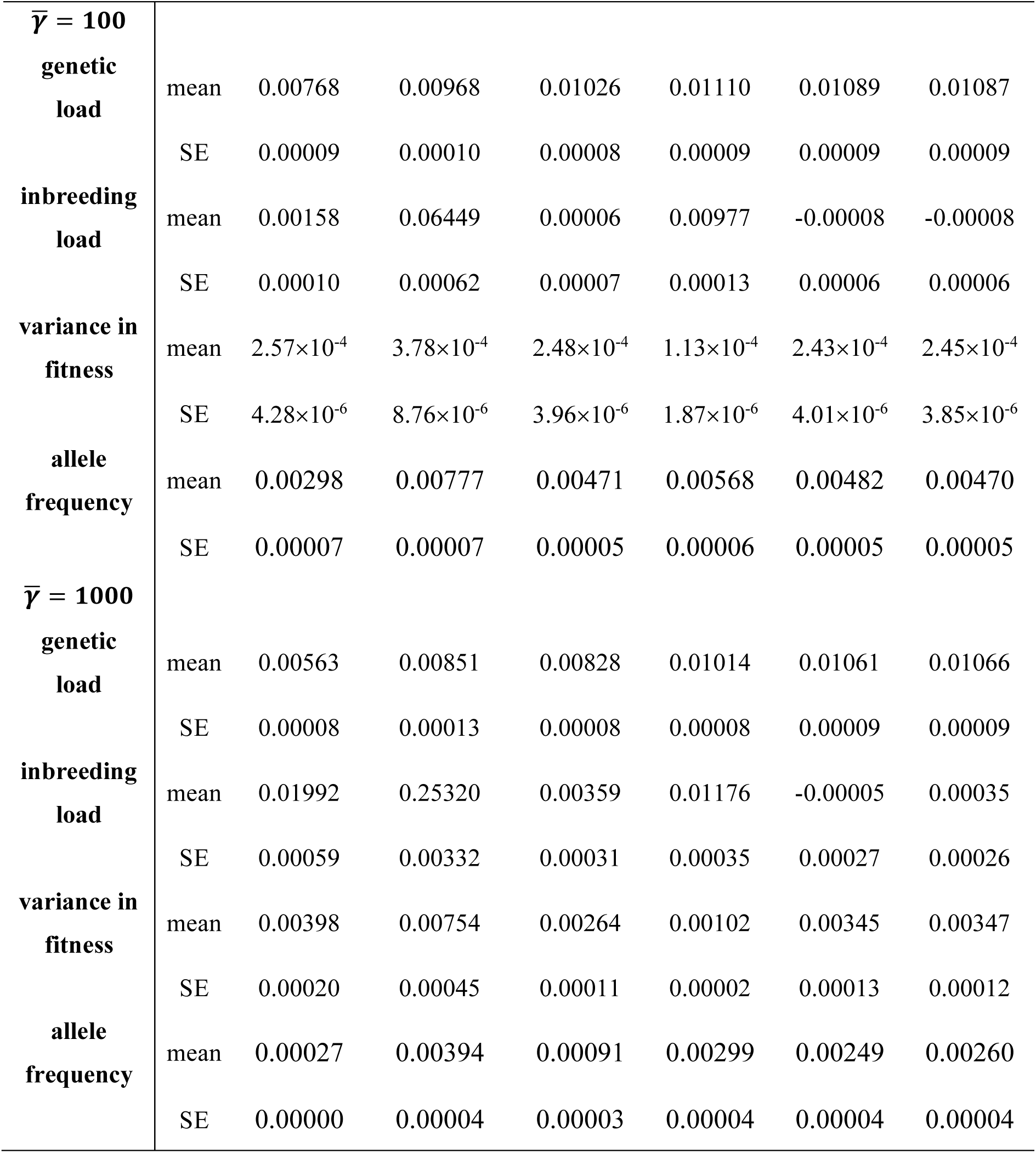
The load statistics under the gene and sites models for a gamma distribution of scaled selection coefficients with shape parameter 0.3 and mean 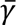 = 2*Ns̅*. 1000 selected sites were simulated, with *Nu* = 0.005 and *Nr* = 0.01, where *N* is the population size (*N* = 1000), *u* is the mutation rate per site/generation, and *r* is the rate of crossing over between adjacent sites. Details of how to estimate the load statistics are given in the Methods section. The means and standard errors (SEs) for 1000 replicate simulations are shown.

The equations for the load statistics derived in section 2 of the Appendix were used to provide the predicted values in Table 3. These are expected to be accurate only for *γ* >> 1; accordingly, it can be seen that the predictions of the genetic and inbreeding loads mostly fit much worse for *γ* = 2 than for *γ* = 20 and *γ* = 100, although the mean frequencies of deleterious alleles are always remarkably close to the predicted equilibrium values when *h* = 0.2 and *h* = 0.5. For the sites model, however, there are strong departures from the predicted equilibrium values with *h* = 0, even for *γ* = 100. This reflects the fact that the mean allele frequency for a fully recessive deleterious mutation in a finite population is expected to be considerably lower than the equilibrium value, as a result of the elimination of deleterious homozygotes produced by drift (Nei 1968). This also causes the inbreeding load for the sites model to be greatly overpredicted when *h* = 0.

For *γ* = 20 and *γ* = 100, the predictions of the values of the genetic and inbreeding loads with *h* = 0.2 and 0.5 are accurate for both the sites and gene models. In contrast, the predicted variances in fitness with *h* = 0 and *h* = 0.2 for these cases are substantially smaller than the variances found in the simulations. The main cause of this difference is likely to be the positive covariance in allelic effects between different sites resulting from LD, which has been ignored in these predictions, although there may also be some contribution from the dominance variance. This effect can be assessed quantitatively, by means of the predicted values of mean *D* at all sites, given by Equations (A19a) of Johri and Charlesworth (2025). Using the formulae of Bulmer (1980, p.158), with equal allelic effects at all selected sites, and assuming that non-additive variance in fitness is negligible, the expected component of variance arising from LD is *C*_*L*_ = 2*G*^2^*D̅V*_*g*_; the net predicted fitness variance is *V* = *V_g_* + *C_L_.* Using the predicted values of *V_g_* in Table 3, this formula yields predictions of 6.85 x 10^−5^ and 6.25 x 10^−5^ for the variances with *γ* = 20 and *h* = 0 and 0.2, respectively; for *γ* = 100, the corresponding values are 1.47 x 10^−4^ and 1.73 x 10^−4^. The corrections are quite substantial for *γ* = 20, and bring the predicted values close to the simulated values, but are negligibly small with *γ* = 100. No contribution from LD is expected for *h* = 0.5, and the predicted variances in fitness match the simulation results in this case. LD is also expected to have minimal effects on *V* for the sites model.

The effect of LD on *V* are counteracted by lower allele frequencies under the gene model, an effect that is especially strong for fully recessive mutations, leading to a much lower *V* under the gene model than the sites model when *h* = 0. However, LD causes *V* to be significantly higher for the gene model than the sites model when *h* = 0.2 and *γ* = 20. With *h* = ½, there is no difference between the two models, as expected. The difference in *V* between the two models thus has a non-linear relation with *h*.

Because the results discussed above are most relevant to *D. melanogaster*-like population parameters, we also conducted simulations with parameters appropriate for human populations (for details, see Johri and Charlesworth 2025), with much smaller values of *Nu* (0.00025) and *Nr* (0.0002). The overall patterns are very similar to those for *Drosophila*-like populations (Table S2 in Supplementary File S1). The mean frequencies of deleterious alleles are significantly smaller with the gene than the sites model; genetic load is slightly lower, while inbreeding load is again drastically smaller for the gene model than the sites model when *h* < ½.

#### Effects of rescaling of population genetic parameters

To evaluate the possible effects of rescaling to keep the products of *N_e_* with the deterministic population genetic parameters such as selection coefficients and mutation rates consistent with their natural population values for *D. melanogaster* (see Dabi and Schrider 2024; Ferrari et al. 2024; Johri and Charlesworth 2025), simulations were also performed with 5000 diploid individuals, where *Nu*, *Nr*, and *Ns* were the same as for *N* = 1000. The mean equilibrium frequencies of deleterious alleles were found to be identical in both cases (compare Table 3 with Table S1 in Supplementary File S1), as is expected from the general theory of population genetic processes when all evolutionary forces are weak and their strength can be represented by single variables such as *u*, *s* and *r* (e.g., Ewens 2004, Chap.5). The genetic and inbreeding loads involve products of the haplotype frequencies and *s*, whereas the variance involves products of the haplotype frequencies and *s*^2^ (see section 1 of the Appendix). If the haplotype frequencies are preserved after multiplying the population size by a factor *C* and keeping the scaled parameters constant, the simulated values of the two load statistics should thus be multiplied by *C* and the variance by *C*^2^, in order to produce values that are comparable with the results for the smaller *N* value (*C* = 5 in the present cast). As can be seen from Tables 3 and S1, the load statistics for *N* = 5000 after this correction agree well with those for *N* = 1000, implying that we can have confidence in the rescaling procedure. These results imply that studies interested in evaluating the extent of mutational load in endangered populations using forward simulations need to apply similar corrections to the load statistics if rescaling is employed at the single gene level, where recombination rate is almost linearly related to map distance.

#### Load statistics with epistasis

With synergistic epistasis between deleterious mutations, a decrease in their mean equilibrium frequencies is expected compared with the case with no epistasis, due to the reduced fitness of genotypes with multiple mutations that is caused by synergistic epistasis (Crow 1970; Kondrashov 1988; Charlesworth 1990) In the simulations with multiple linked sites under a DFE with a relatively large mean scaled selection coefficient (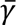 = 2*Ns̅* = 100) and varying degrees of epistasis (with *ε* ranging from 0.02 to 0.32), there is a severe decrease in the mean equilibrium frequency of deleterious mutations with increasing *ε*, for both the gene and sites models (Figure 1). Even with the lowest strength of epistasis (*ε* = 0.02), there is no significant difference in mean equilibrium frequencies of deleterious alleles between the gene and sites models. There is a corresponding decrease in the inbreeding load (*B*) with increasing *ε* for both the gene and sites model (Figure 2) when *h* < ½.

**Figure 1:**
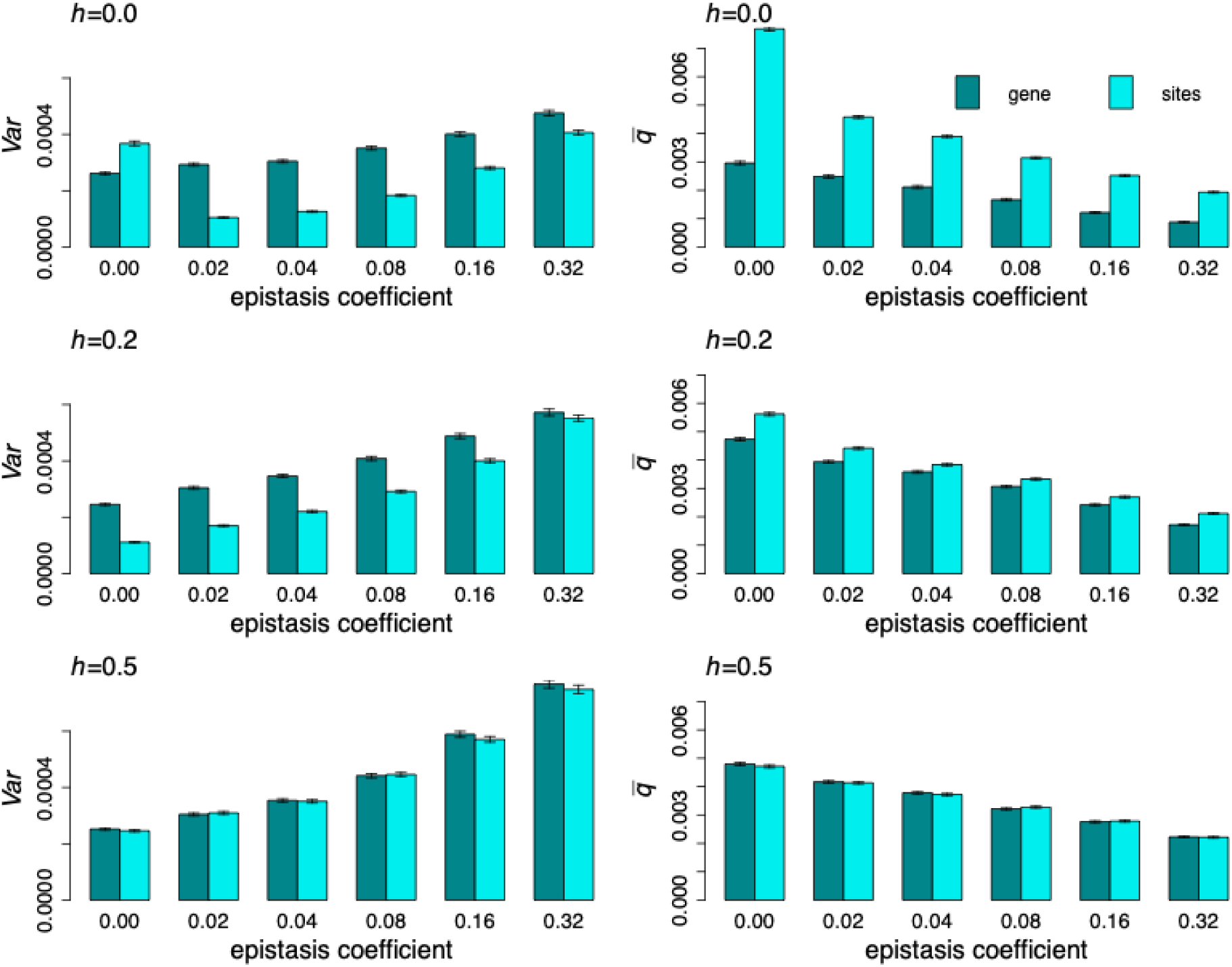
The variance in fitness (*Var*) and mean deleterious allele frequency (*q̅*) for the gene and sites models with the *Drosophila*-like simulation parameters, and varying values of the epistasis coefficient (*ε*). Scaled selection coefficients followed a gamma distribution with 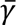 = 100 and shape parameter 0.3. 1000 selected sites were simulated with *Nu* = 0.005 and *Nr* = 0.01, where *N* is the population size (*N* = 1000), *u* is the mutation rate per site and *r* is the rate of crossing over between adjacent sites. The means and standard errors (SEs) for 1000 replicate simulations are shown.

**Figure 2:**
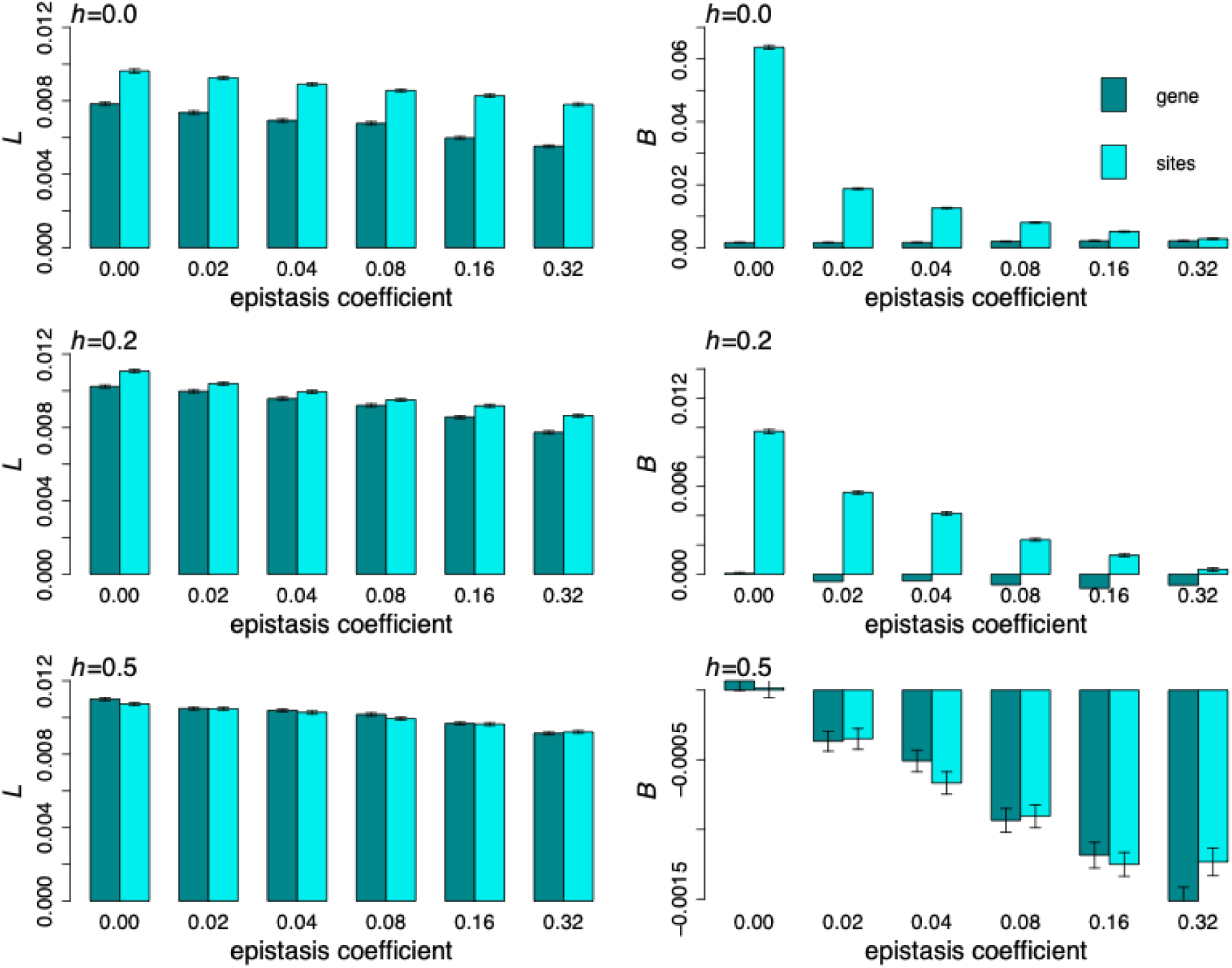
The means across replicate simulations of the genetic load (*L*) and inbreeding load (*B*) for the gene and sites fitness models, with varying values of the epistasis coefficient (*ε*). The selection coefficients follow a gamma distribution with 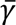 = 100 and shape parameter 0.3. 1000 selected sites were simulated, with *Nu* = 0.005 and *Nr* = 0.01, where *N* is the population size (*N* = 1000), *u* is the mutation rate per site/generation, and *r* is the rate of crossing over between adjacent sites. The error bars represent the SEs across 1000 replicate simulations.

*B* is still significantly and drastically higher with the sites model than the gene model for both *h* = 0 and *h* = 0.2. There is no difference between the two models with *h* = 0.5, as expected; unexpectedly, however, *B* becomes increasingly slightly negative as *ε* increases. One possible explanation is that this is an artefact of the way *B* in which was estimated; the fitness of inbred individuals was calculated by sampling 100 haploid genomes, whereas the mean fitness was calculated using the entire population. However, even if we calculate the fitness of outbred individuals by sampling 100 diploid individuals, *B* remains negative for these parameter sets (see Figure S1 in Supplementary File S1), so that this explanation can be ruled out.

The probable cause of this effect is as follows. With the parameter set in question, the frequencies of deleterious alleles are sufficiently high that essentially all haplotypes carry at least one deleterious mutation. In the two-site case with *h* = 0.5, a – +/+ – individual has fitness reduction *s*(1 + *ε*), whereas – + or + – homozygotes have a fitness reduction *s* (there is no effect of epistasis). Thus, the heterozygote has a lower fitness than the homozygote. In other words, the outbred population has more opportunities for pairwise fitness interactions that reduce fitness than the inbred population, resulting in slightly negative inbreeding depression. Epistatic interactions of this kind have been studied theoretically and empirically in connection with the biometrical genetics of crosses between inbred lines (e.g., Wright 1968, Chap. 15, pp. 391-403; Jinks 1983). A lower value of a fitness component in the F_1_ compared with the mean of the two parents of the kind expected under this model appears to be uncommon in such crosses, with a prevalence of the opposite pattern (heterosis), consistent with the effects of largely recessive effects of deleterious alleles or heterozygote advantage outweighing epistatic effects of this kind.

There is a much smaller effect of epistasis on the genetic load (*L* in Figure 2), with a minor and steady decrease in *L* with an increase in *ε* for both fitness models. As before, the difference in the genetic load between the two models is relatively small and only significant when mutations are completely recessive. The variance in fitness (*V*) also increases with an increase in the epistasis coefficient when *h* = 0.2 or 0.5 (and only very slightly when mutations are fully recessive). However, as was also found without epistasis, the variance is much larger for the gene model compared to the sites model, especially when mutations have increased recessivity and intermediate/low levels of epistasis (Figure 1). With high levels of epistasis (*ε* = 0.32), however, there is no difference in *V* between the two fitness models. In summary, if there were widespread epistasis between deleterious mutations, there would be no substantial difference in mean equilibrium allele frequencies between the two models, but the inbreeding and genetic loads would be lower with the gene model than the sites model if mutations have moderate to high recessivity. Note that with much lower rates of crossing over (0.1 × the standard rate used here), the behavior of the load statistics remains the same as described above (Figures S1 and S2 of Supplementary File S1).

## Discussion

### Load statistics with no epistasis

#### Differences between the gene and sites models

The main conclusion from the analytical and simulation results for the case of no epistasis is that the inbreeding load *B* is always much smaller under the gene model than the sites model, being effectively zero for 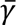 ≤ 20, unless the dominance coefficient *h* is close to 0.5; in this case *B* is close to zero for both models, as expected from basic theory (Morton, Crow and Muller 1956). A similar pattern is seen with synergistic epistasis, but *B* becomes very small for both the gene and sites models, and even slightly negative, when epistasis is strong (Figure 2). These results raise the question of why *B* should be so much smaller under the gene model than the sites model. One factor is the smaller mean frequency of deleterious alleles (*q̅*) for the gene model than the sites model, at least when *γ* >> 1. However, the relation of *q̅* to the strength of selection does not track the behavior of *B*. For example, for the case of a gamma distribution with *h* = 0.2, *B* and *q̅* for the gene model with 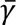 = 100 are approximately 6% and 30% of the values for the sites model, respectively; when 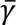 = 1000, *B* and *q̅* for the gene model are 30% and 12% of the sites model values, respectively (see Table 4).

This discrepancy probably arises from the fact that haplotypes carrying deleterious mutant alleles can be relatively common when the number of selected sites in a gene (*G*) is large. For example, with *γ* = 20 and *h* = 0.2 the equilibrium frequency of deleterious alleles is approximately 0.001 (see Table 3) so that the mean number of mutant alleles per haploid genome with *G* =1000 is 1000 x 0.001 = 0.1. Genotypes with deleterious mutations in *trans* are thus relatively common, so that a heterozygous mutation at a given site behaves more like a dominant mutation than under the sites model, reducing its contribution to inbreeding depression. It is surprising that such a large difference in the expectation for the level of inbreeding depression caused by a single gene can be produced by the lack of complementation between mutations in *trans*.

Another way of looking at this effect is to note that *B* for the additive gene model when *γ* >> 1 is approximately (1 − 2*h*)*μe*^−*μ*^*s* (see Equation A8*b*), where *μ* = *Gq̂* is the mean number of mutations per haplotype and *q̂* is the deterministic equilibrium frequency of mutant alleles. In contrast, *B* for the additive sites model in this case is equal to *Gu*(1– 2*h*)/*h* when *h* > 0 (Morton et al. 1956; Charlesworth and Hughes 2000). When *h* > 0 and *μ* << *h*, Equation (A21*b*) of Johri and Charlesworth (2025) shows that *q̂* approaches the sites value of *u*/(*hs*). The values of *B* for the two models thus converge when selection is sufficiently strong, and *G* is sufficiently small that *e*^−*μ*^ ≈ 1. A similar result holds when *h* = 0.

There is a lack of information on whether or not the lack of complementation assumed in the gene model applies to deleterious mutations with minor effects on fitness, as opposed to the loss-of-function mutations that have generally been studied in relation to complementation. There are, however, a number of studies of the phenotypes of heterozygotes for hypomorphic mutations in the same gene, suggesting that partial impairment of functionality is usually associated with lack of complementation. Wright (1968, p.70) gives example of multiple alleles at coat color loci of the guinea pig that show this behavior, and Chandler et al. (2017) describe similar effects for two wing size loci of *D. melanogaster*. Quantitative studies of dominance effects on fitness in diploids such as yeast (*e*.*g*., Matsui et al. 2022), with a focus on measuring complementation, are needed to shed further light on this question.

#### Other models of deleterious mutations

We now consider the relation of the gene model to the widely used model that treats a gene as a single non-recombining unit subject to mutation from wild-type alleles to mutant alleles, each of which causes a fitness reduction of *s* when homozygous or in combination with each other, and *hs* when heterozygous with wild-type (e.g. Charlesworth et al. 1992; Wang et al. 1999; Kamran-Disfani and Agrawal 2014; Bersabé et al. 2016). This type of model is equivalent to a single locus with mutation at rate *Gu* from wild-type to a mutant allele with selection coefficient *s*.

At first sight, this situation seems equivalent to the gene model with identical selection coefficients for each mutation. However, with the relatively weak selection against many deleterious mutations suggested by population genomics data (see section 1 of Supplementary File S2), the ratio of *Gu* to *hs* is likely to be such that most haplotypes segregating in the population carry at least one mutation. Even with relatively strong selection (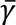 =1000 and *h* = 0.2 in Table 4), the mean number of deleterious mutations per haploid genome for a sequence of 1000 selected sites is 1000 x 0.00091= 0.91.

The assumption that all deleterious mutations are only one step away from wild-type may thus be unsafe. This has important consequences for the predicted values of the genetic and inbreeding loads. Consider the case of a fixed selection coefficient with *γ* = 20 described in Table 3. In this case, the mean deleterious allele frequency per site is 0.00098 under the gene model, with a very small standard error. With the weak LD associated with these parameters (see Figure 3 of Johri and Charlesworth 2025), the distribution of the number of mutations per haploid genome closely follows a Poisson distribution, so that (disregarding variability between replicate simulations) the probability of a mutant-free haplotype is approximately exp(– 0.98) ≈ 0.38. The predicted frequency of haplotypes containing at least one mutation is thus 0.62. If this is used as the deleterious allele frequency *q* in the standard formulae for *L* and *B* (*L* = [2*h* + (1 − 2*h*)*q*]*sq*; *B* = *sq* − *L*), we obtain *L* = 0.0048 and *B* = 0.0014. These are substantially lower than the predicted values of *L* = 0.0078 and *B* = 0.0022 under the gene model, reflecting the fact that this calculation ignores the contributions of haplotypes carrying more than one mutation. One-step mutational models can therefore seriously underpredict the load statistics, producing a bias in the opposite direction to that described in the previous section.

#### Implications for data on inbreeding load

Recent studies of genetic and inbreeding loads that have used population genomics data to simulate populations subject to deleterious mutations and to predict genetic and inbreeding loads (reviewed by Kyriazis, Robinson and Lohmueller 2023) have mostly used the sites model. Multiplicative fitnesses across selected sites and genes have also usually been assumed in models of inbreeding load – but see Charlesworth (1998) and García-Dominguez et al. (2018) for exceptions. Our results are for a single gene, so it is important to consider what happens if they are extrapolated to the whole genome. For convenience, we will assume purely autosomal inheritance, leading to a probable overestimation of the load statistics for the entire genome, given that the X chromosome is associated with lower loads than autosomes due to selection against hemizygous mutations in males (Wilton and Sved 1979).

To scale up to multiple genes, we assume that the sites model applies to between-gene effects for both models of single genes, due to complementation between non-allelic mutations. As in previous studies, starting with Haldane (1937), we assume multiplicative fitness effects across different genes; with small selective effects the differences from additive effects will be small in most cases. Our simulation results were based on a scaling of population size from 10^6^ to 10^3^; to generate realistic results for a *Drosophila*-like system, we would need to rescale back to the *N* = 10^6^ values; this means dividing linear quantities in *s* by 10^3^ and quadratic quantities such as the variance by 10^6^ (see the discussion of rescaling in the section *Load statistics with no epistasis*). If we assume 14000 genes in the *Drosophila* genome (Misra et al. 2002), the individual gene linear quantities (the loads and inbreeding loads) in the tables should thus be multiplied by 14 and the variance by 0.014. This ignores any contributions from mutations in non-coding sequences, which will be considered below.

In general, let the linear and quadratic multipliers reflecting rescaling be *f* and *f*^2^, and the number of genes be *n*. For the inbreeding load, defined as the difference between the natural logarithms of outbred mean fitness and fully inbred mean fitness, the *B* values in the tables need only be multiplied by *nf* to obtain the total inbreeding load, *B_T_*. If the mean load *L* for each gene is small, the total genetic load with multiplicative effects across *n* genes is given by the following expression:

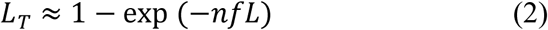

Similarly, if there is no LD between different genes, using the rule that the expectation of a product of a set of independent variables is the product of their expectations, the total variance with multiplicative effects across genes is given by:

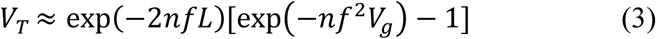

Given *n* and *f*, the values of the relevant statistics in the tables can easily be converted into values for the whole genome by using these relations. Using the mean values of the load statistics in Table 4 with 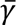 = 1000, Table 5 shows the values obtained with *n* = 14000 and *f* = 14; *σ_T_* is the standard deviation corresponding to *V_T_*. These results show that the difference between the two models has only a small effect on the expected genome-wide genetic load and genetic variance/standard deviation (mean *s* is only 5 x 10^−4^ after rescaling *N*), but that the gene model has a drastically smaller total inbreeding load than the sites model with *h* = 0.2 (0.050 versus 0.165), despite the fact that the sites model applies to between-locus fitness effects.

**Table 5.**
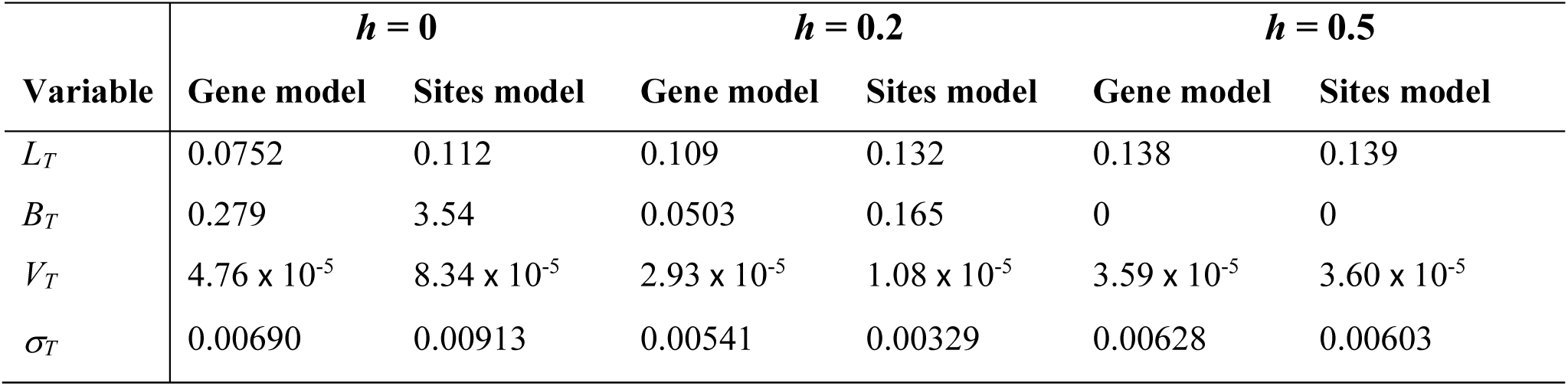
The values of the load statistics obtained from Table 4 with 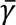 = 1000 when scaled up to the whole *D. melanogaster* genome.

Our results suggest that the sites model of mutation and selection is likely to considerably *overpredict* the value of *B*, unless the mean strength of selection is so large and the distribution of selection coefficients is so tight that the population can be treated as fully deterministic. Population genomics studies of the DFE for nonsynonymous mutations in organisms such as humans (Kim et al. 2017) and *Drosophila melanogaster* (Campos et al. 2017; Johri et al. 2020) suggest that a substantial fraction of deleterious mutations are subject to the joint effects of drift and selection (see section 1 of Supplementary File S2).

This conclusion is important, because reconciling observed and predicted values of *B* has been the subject of much debate, with some workers asserting that the observed values can be explained by mutation-selection balance with multiplicative fitnesses (e.g., Pérez-Pereira et al. 2021; Robinson et al. 2023), while others have proposed that additional factors (such as variation due to balancing selection) are likely to be involved (e.g., Charlesworth 2015). If it can be shown that empirical estimates of *B* are incompatible with values predicted on the basis of the sites model, using well-grounded estimates of mutation rates, dominance coefficients and realistic demographic histories, then contributions from balancing selection or synergistic epistasis of the type modeled by Charlesworth (1998) must be involved.

A difficulty with this approach is that the currently used methods for inferring the DFE from population genomic data assume the sites model for predicting variant frequency distributions at sites under selection. As pointed out in section 1 of Supplementary File S2, this means that the mean strength of selection as measured by 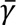 is likely to be overestimated, given that the gene model predicts lower deleterious variant frequencies than the sites model. Further research into the magnitude of this problem needs to be carried out, but the fact that it is likely to inflate the predicted sizes of the genetic and inbreeding loads needs to be borne in mind. Moreover, interpreting empirical estimates of the extent of inbreeding is problematic because reliable estimates of *B* for net fitness in natural populations are hard to obtain, with substantial disagreements among different workers concerning the typical magnitude of *B* (Robinson et al. 2023). This makes it difficult to compare the data on inbreeding load with theoretical predictions.

#### Inbreeding load in Drosophila

In order to avoid the problems associated with interpreting measurements of fitness in wild populations, we here compare theoretical estimates of levels of inbreeding load with estimates obtained from laboratory experiments on *D. melanogaster*. An ingenious method for estimating *B* for net fitness under laboratory conditions for a single major chromosome of some *Drosophila* species was devised by John Sved, known as the balancer equilibration (BE) technique (Sved and Ayala 1970). This is based on using population cages to compete wild-type chromosomes extracted from natural populations (chosen to be homozygous viable and fertile) against homozygous lethal balancer chromosomes that suppress crossing over when heterozygous. The equilibrium frequencies of wild-type chromosomes can be used to estimate the ratio *R* of the mean fitness of chromosomal homozygotes to the mean fitness of chromosomal heterozygotes (reviewed by Frankham 2023). The mean value of *R* for chromosome 2 of *D. melanogaster* over two well-replicated experiments was 0.18±0.03, giving an estimate of *B* = – ln(*R*) of 1.7 (Frankham 2023). The BE results on the data for all three major chromosomes imply that *B* for the whole genome of *D. melanogaster* (including the contribution from homozygous lethal mutations) is approximately 5 (Frankham 2023).

The analysis of population genetic data for *D. melanogaster* presented in section 3 of the Appendix yields a prediction of *R* = 0.66 and *B* = 0.42 for chromosome 2 under the sites model. Here, possible effects of purifying selection at non-coding sites as well nonsynonymous sites and strongly selected (but non-lethal) mutations are taken into account, using fairly liberal assumptions with respect to the magnitude of *B*, so that the resulting values are likely to overestimate *B*. This value of *B* uses the deterministic expressions for *R* and *B* given by Equations (A9) and (A10), and assumes an unrealistically low *h* value of 0.125. In addition, an approximate correction for the effects of drift was obtained by multiplying *u* by the proportion of mutations with 2*N_e_hs* < 2.5 under a gamma distribution with shape parameter 0.3, thereby removing the contribution of nearly neutral mutations (see section3 of the Appendix).

Given that the gene model predicts a substantially lower value of *B* than the sites model, *B* = 0.42 for chromosome 2 is almost certainly an overestimate, but is still less than 25% of the observed value. There thus appears to be much more inbreeding load due to deleterious mutations under laboratory conditions in *D. melanogaster* than can be accounted for purifying selection under a multiplicative fitness model, especially considering the various biases towards overpredicting the inbreeding load caused by deleterious mutations.

#### Why is the inbreeding load reduced when N_e_s is small?

Tables 3 and 4 show that, other things being equal, smaller values of *B* are associated with smaller values of *N_e_s*. Numerical methods for obtaining an exact single-locus prediction for *B* in a finite population have been developed by Charlesworth (2018), and extensive multi-locus simulation results using a variety of parameter values have been described in the literature (Kyriazis et al. 2023). Both of these approaches show that the level of inbreeding load caused by deleterious mutations is positively related to *N_e_s* under a wide range of model assumptions. As has been widely discussed in the literature (see review by Robinson et al. 2023), there are two possible reasons for this effect – the “purging” of deleterious, strongly recessive variants from small populations as a result of their exposure to selection against homozygotes (Nei 1968), and the loss of variability, including the (temporary) fixation of deleterious variants (Kimura et al. 1963). The analysis presented in sections 2 and 3 of Supplementary File S2 shows that the second factor is responsible for the effects seen in our simulations.

The weak effects of drift on the genetic load and variance in fitness seen in Table 4 are at first sight surprising, given that with 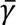 = 2 a substantial proportion of the distribution of selection coefficients falls below the nearly neutral threshold of *hγ* < 2.5 described above, where large departures from the deterministic values are expected. However, this part of the distribution involves very small selection coefficients, and thus makes only a small contribution to the genetic load and the variance in fitness. This mitigates the effect of drift in causing an increase in load. In contrast, with a fixed scaled selection coefficient of *γ* = 2, there is a substantial increase in load compared with the higher value of *γ* = 20 (Table 3).

### The role of epistasis

#### Epistasis and inbreeding load

One important aspect of our simulation results with synergistic epistasis is that *B* for the gene and sites models is strongly reduced by even a small amount of epistasis when *h* < ½, despite only a small reduction in load; there is a much smaller effect on *B* under the gene model, where it is always small (Figure 2). This reduction in *B* with increasing epistasis is in sharp contrast to the model of Charlesworth (1998), which predicts that synergistic epistasis causes an *increase* in *B* when the logarithm of fitness is a quadratic, decreasing function of the number of deleterious mutations. As shown in section 4 of the Appendix, when only homozygous genotypes are considered, the pairwise epistasis model used here is identical to the quadratic model. The opposite behavior of the two models is thus surprising at first sight.

The explanation lies in the fact that the model of Charlesworth (1998) assumes that *h* > 0 and the quadratic term for heterozygous mutations is proportional to the negative of the product of *h*^2^ and the square of the number of heterozygous mutations (*n*^2^). In contrast, the pairwise epistasis model assumes that this term is proportional to – *hn*^2^. This means that the fitness of the outbred population is relatively insensitive to the quadratic fitness term under the first model, since most mutations are present in heterozygotes. However, the fully inbred genotypes have strongly reduced fitnesses due to the quadratic term, resulting in an increase in *B*. Under the pairwise model, there is a sharp decline in *q̅* with increasing *ε*, while the load is relatively unaffected (compare Figures 1 and 2). Since *B* is strongly determined by *q̅*, the net result is that synergistic epistasis reduces *B*, even when *h* = 0.

The consequences of epistasis for inbreeding depression are thus highly model dependent, and it cannot be assumed that synergistic epistasis necessarily leads to increased inbreeding depression compared with the multiplicative fitness model, as has been done in the past (Crow 1970; Charlesworth 1998; García-Dominguez et al. 2018). The pairwise model has the advantage of greater flexibility in modeling the fitnesses of heterozygotes and differences in selection coefficients between sites, whereas the quadratic model is forced to make somewhat arbitrary assumptions about the effects of dominance (for details, see Charlesworth 1998). However, it is important to note that we have only developed this model for within-gene interactions, and for pairwise interactions within genes; the nature of epistatic interactions between deleterious mutations in different genes and their consequences for the load statistics, as well as the effects of higher order interactions, remain to be explored.

### Conclusions

Our results show that the gene model predicts much lower levels of inbreeding depression than are expected from current mutational models, and that a revised model of synergistic epistasis between mutations within genes leads to even smaller values of inbreeding depression, making it hard to explain observed values of the inbreeding load on the purely mutational hypothesis. It should, of course, be borne in mind that complex demography, including strong bottlenecks and hidden population structure, may also play a role in the observed mismatch between expected and observed estimates of *B*. More organism-specific simulations that take demographic history into account, as well as the differences between the gene and sites models, are likely needed to understand the respective contributions of deleterious mutations and balancing selection to inbreeding depression.

We note that much of the current literature on inbreeding depression assumes that this is caused almost entirely by deleterious mutations (e.g., Bertorelli et al. 2022; Peréz-Pereira et al., 2022; Robinson et al. 2023), despite evidence to the contrary dating back over many years (e.g., Charlesworth and Hughes 2000; Charlesworth 2015). The contribution of balancing selection to inbreeding depression should thus be further examined (*e.g.*, González-Castellano et al. (2025).

## Acknowledgments

We thank Kirk Lohmueller, Denis Roze and three anonymous reviewers for their helpful comments on an earlier version of the manuscript. This research was conducted using computational resources provided by ITS Research Computing at the University of North Carolina at Chapel Hill. PJ was funded by the National Institute of General Medical Sciences of the National Institutes of Health under the award number R35GM154969. We

## Appendix

### The load statistics for the two-site deterministic models

Using the fitness model of Table 1, and the notation for the haplotype frequencies described in the main text, the genetic load under the gene model is given by:

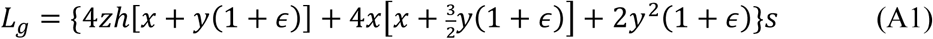

The inbreeding load is:

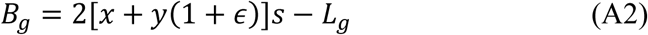

and the variance in fitness is:

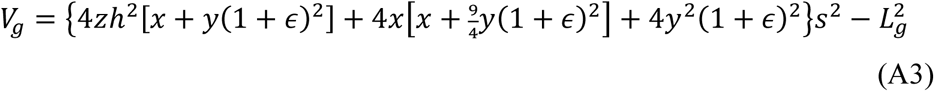

Under the sites model, the corresponding formulae are:

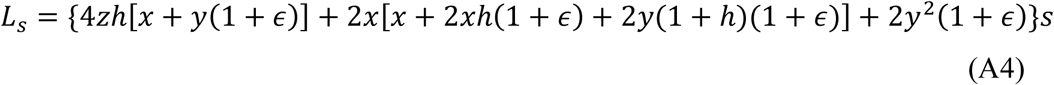

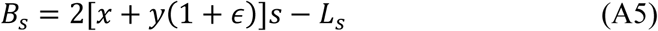

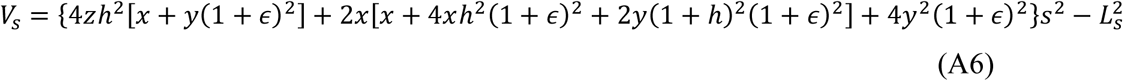

In the case of complete recessivity, Equations (A1) and (A4) for the genetic loads under the gene and sites models without epistasis reduce to the following expressions:

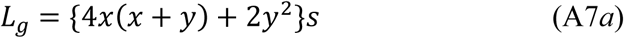

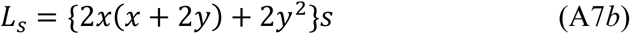

Since *y* << *x*, the dominant terms in these equations are 4*x*^2^*s* and 2*x*^2^*s*, respectively. It follows that the load is higher for the gene than the sites mode when *h* = 0. By continuity, this will also be true when *h* is sufficiently small, as can be seen in Table 2 for the case of *h* = 0.01.

### The load statistics for the multi-site deterministic models without epistasis

The formulae for the additive multi-site gene model derived in the last section of the Appendix of Johri and Charlesworth (2025) can be used to obtain expressions for the load statistics for the case of a large number (*G*) of sites with the same *s* and *h*, when *N_e_s* is sufficiently large that deterministic results should apply. We first note that the equilibrium frequency of a deleterious mutation at a given site (*q̂*) can be found using numerical iteration of Equation (A21*a*) of Johri and Charlesworth (2025), with the mean number of deleterious mutations per haploid gene given by *μ* = *Gq̂*. Under the assumptions used to derive this equation (including the condition *q̂* ≪ 1), the equilibrium genetic load for the gene model is given by the following expression:

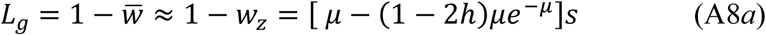

The inbreeding load is given by:

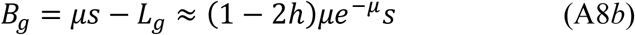

The variance in fitness contributed by the sum of the variances at individual sites (the genic variance) is given by:

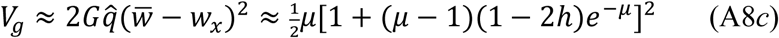

The corresponding expressions for the sites model can be found from the products of *G* with the standard single-locus expressions for deterministic mutation-selection equilibrium when *q̂* << 1 (see Charlesworth and Hughes 2000), on the assumption that LD is negligible. In this case, 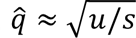 when *h* = 0, and *q̂* ≈ *u*/(*hs*) when *h* is bounded away from zero (Haldane 1937).

### Relating population genomics data to the classical expression for inbreeding depression

Here we attempt to relate inferences from population genomics data on *D. melanogaster* to the empirical results on the effect on fitness of homozygosity for chromosome 2 of this species, described in the section of the Discussion *Inbreeding load in Drosophila.* The ratio of the mean fitnesses of chromosomal homozygotes and heterozygotes under the multiplicative fitness model with deterministic mutation selection balance for mutations with *h* > 0 is:

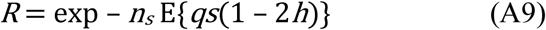

where *n_s_* is the mean number of sites on the chromosome that are subject to significant selection (provisionally identified as nonsynonymous sites in coding sequences), *q* is the frequency of a deleterious variant at a given nonsynonymous (NS) site, *s* and *h* are the selection and dominance coefficients, and E{} denotes the expectation over all NS sites.

We have *q* ≈ *u*/(*hs*) for strongly selected mutations, so that:

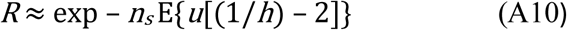

This is the exponential of the negative of the inbreeding load (*B*), under mutation-selection balance with partial recessivity, assuming multiplicative fitnesses across loci (Morton et al. 1956). This formula applies even when there are significant effects of drift on *q*, provided that 2*N_e_hs* is sufficiently large and mating is random, since the distribution of *q* then follows a gamma distribution with mean *u*/(*hs*) and variance *u*/[4*N_e_*(*hs*)^2^] (Nei 1968).

In section 1 of Supplementary File S2, we present an analysis of relevant population genomics data on *Drosophila melanogaster,* where it was proposed that 91% of NS sites are under significant selection (such that 2*N_e_hs* ≥ 2.5); it can be assumed that the remainder are nearly neutral, with negligible effects on fitness. Given that chromosome 2 is 37% of the genome, that there are approximately 14000 genes in the fly genome (Misra et al. 2002). Assuming a mean of 1500 exonic sites per gene and that 70% of exonic sites are NS (Campos and Charlesworth 2019), *n_s_* for chromosome 2 is 0.37 x 0.91 x 14000 x 0.70 x 1500 = 4.95 x 10^6^. With the mutation rate of *u* = 4 x 10^−9^ proposed in section 1 of Supplementary File S2, and the value of *h* = 0.25 suggested by Manna *et al*. (2011), this gives *R* = exp – 0.0396 = 0.961. If we multiply *n_s_* by three, to include selection on non-coding sites, *R* = 0.888. The strength of selection is irrelevant to this calculation, so that the mean of 2*N_e_hs* could be very different from that used above but the value of *R* would be the same.

Variation in *h* around 0.25 would, of course, increase these estimates. If there were a two-fold increase in 1/*h*, corresponding approximately to a squared coefficient of variation of *h* of 1 (as under an exponential distribution), or to a fixed value of *h* = 0.125, then *B* = – ln *R* would be increased to three times that corresponding to the above value, giving *B* = 0.356, *R* = 0.700 (this is likely to be an underestimate, since small values of *h* will reduce the proportion of significantly selected mutations). If the above estimate of the contribution from major effect mutations is also included, *R* becomes 0.656, *B* = 0.421.

The remaining question is whether nearly neutral sites, which are ignored in these calculations, contribute significantly to inbreeding depression. As just described, the effect of drift away from mutation-selection equilibrium is to reduce *B*, so that use of the deterministic formula will overestimate it for nearly neutral sites. If 9% of new mutations fall into this category, their maximal contribution to *B* would be 0.09/0.91 = 0.10 of the values calculated above, a fairly trivial amount.

### Relating two models of epistasis

A widely used model of epistasis among deleterious mutations assumes that either fitness or the logarithm of fitness has a quadratic relation to the number of deleterious mutations (Crow 1970; Charlesworth 1990, 1998). This raises the question of its relation to the pairwise interaction model used in the present paper, which we now examine. For simplicity, consider the fitness of homozygotes only, with fixed selective effects across sites. Under the quadratic model, the fitness of an individual carrying *n* mutations can be described by a function of the following form:

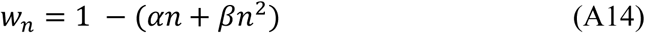

Under the pairwise model, we have:

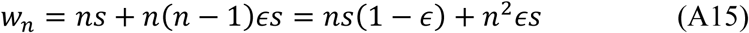

Equating the coefficients of *n* and *n*^2^ in these two equations gives the relations

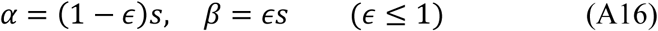

This result implies that the quadratic model inherently involves pairwise epistasis, at least in its simplest realization, when only homozygotes are considered. The pairwise model has the advantage of greater flexibility in modeling the fitnesses of heterozygotes and differences in selection coefficients between sites, whereas the quadratic model is forced to make somewhat arbitrary assumptions about the effects of dominance (Charlesworth 1998).

