## Supplementary File S1 for "A gene-based model of fitness and its implications for genetic variation: Genetic and inbreeding loads"

### FILE S1. SUPPLEMENTARY FIGURES AND TABLES

**Table S1:** The load statistics under the gene and sites models when a larger than standard Wright-Fisher population of 5000 diploid individuals relevant to *Drosophila*-like populations was simulated. Scaled selection coefficients followed a gamma distribution with shape parameter 0.3 and mean $\bar{\gamma} = 2N\bar{s} = 100$. 1000 selected sites were simulated, with $Nu$ = 0.005 and $Nr$ = 0.01, where $N$ is the population size ($N$ = 5000), $u$ is the mutation rate per site/generation, and $r$ is the rate of crossing over between adjacent sites. Details of how to estimate the load statistics are given in the Methods section. The means and standard errors (SEs) for 1000 replicate simulations are shown. Note that "observed directly" refers to the load statistics estimated directly from the simulations (as given in the Methods section). In order to compare the load statistics to Table 4, a correction for rescaling is required as follows: the means and SEs of genetic and inbreeding loads were multiplied by 5 while those of the variance in fitness were multiplied by 25 (see the main text for an explanation).

| | | ***h*=0.0** | | ***h*=0.2** | | ***h*=0.5** | |
|---|---|---|---|---|---|---|---|
| | | **Gene model** | **Sites model** | **Gene model** | **Sites model** | **Gene model** | **Sites model** |
| **genetic load** | *observed directly* | | | | | | |
| | mean | 0.00151 | 0.00193 | 0.00205 | 0.00222 | 0.00221 | 0.00218 |
| | SE | 0.00002 | 0.00002 | 0.00002 | 0.00002 | 0.00002 | 0.00002 |
| | *corrected for rescaling* | | | | | | |
| | mean | 0.00756 | 0.00963 | 0.01023 | 0.01108 | 0.01106 | 0.01091 |
| | SE | 0.00009 | 0.00010 | 0.00008 | 0.00009 | 0.00009 | 0.00009 |
| **inbreeding load** | *observed directly* | | | | | | |
| | mean | 0.00035 | 0.01279 | 0.00001 | 0.00194 | 0.00001 | 0.00002 |
| | SE | 0.00002 | 0.00012 | 0.00001 | 0.00003 | 0.00002 | 0.00002 |
| | *corrected for rescaling* | | | | | | |
| | mean | 0.00177 | 0.06393 | 0.00005 | 0.00970 | 0.00005 | 0.00008 |
| | SE | 0.00011 | 0.00061 | 0.00007 | 0.00013 | 0.00008 | 0.00009 |
| **variance in fitness** | *observed directly* | | | | | | |
| | mean | $1.02\times10^{-5}$ | $1.52\times10^{-5}$ | $9.92\times10^{-6}$ | $4.44\times10^{-6}$ | $1.02\times10^{-5}$ | $1.00\times10^{-5}$ |

| | | | | | | | |
|---|---|---|---|---|---|---|---|
| | SE | $1.63\times10^{-7}$ | $2.99\times10^{-7}$ | $1.46\times10^{-7}$ | $6.85\times10^{-8}$ | $1.53\times10^{-7}$ | $1.47\times10^{-7}$ |
| | *corrected for rescaling* | | | | | | |
| | mean | $2.56\times10^{-4}$ | $3.80\times10^{-4}$ | $2.48\times10^{-4}$ | $1.11\times10^{-4}$ | $2.55\times10^{-4}$ | $2.51\times10^{-4}$ |
| | SE | $4.07\times10^{-6}$ | $7.48\times10^{-6}$ | $3.64\times10^{-6}$ | $1.71\times10^{-6}$ | $3.82\times10^{-6}$ | $3.69\times10^{-6}$ |
| **allele frequency** | mean | 0.00285 | 0.00764 | 0.00460 | 0.00572 | 0.00481 | 0.00475 |
| | SE | 0.00006 | 0.00007 | 0.00005 | 0.00005 | 0.00005 | 0.00005 |

**Table S2.** The load statistics for the gene and sites models in human-like populations. 2000 selected sites were simulated with $Nu$ = 0.00025 and $Nr$ = 0.0002, where $N$ is the population size ($N$ = 1000), $u$ is the mutation rate per site/generation, and $r$ is the rate of crossing over between adjacent sites. Selection coefficients followed a gamma distribution with a shape parameter of 0.23 and mean $\bar{\gamma} = 2N\bar{s} = 850$. Details of how to estimate the load statistics are given in the Methods section. The means and standard errors (SEs) for 1000 replicate simulations are shown.

| | | ***h*=0.0** | | ***h*=0.2** | | ***h*=0.5** | |
|---|---|---|---|---|---|---|---|
| | | **Gene model** | **Sites model** | **Gene model** | **Sites model** | **Gene model** | **Sites model** |
| **genetic load** | mean | 0.00061 | 0.00070 | 0.00090 | 0.00097 | 0.00096 | 0.00096 |
| | SE | 0.00002 | 0.00003 | 0.00002 | 0.00003 | 0.00003 | 0.00003 |
| **inbreeding load** | mean | 0.00527 | 0.02158 | 0.00056 | 0.00094 | 0.00004 | 0.00000 |
| | SE | 0.00038 | 0.00092 | 0.00011 | 0.00009 | 0.00007 | 0.00008 |
| **variance in fitness** | mean | 0.00030 | 0.00039 | 0.00021 | 0.00009 | 0.00020 | 0.00023 |
| | SE | 0.00004 | 0.00007 | 0.00003 | 0.00001 | 0.00002 | 0.00002 |
| **allele frequency** | mean | 0.00010 | 0.00028 | 0.00018 | 0.00024 | 0.00020 | 0.00020 |
| | SE | 0.00000 | 0.00001 | 0.00001 | 0.00001 | 0.00001 | 0.00001 |

**Figure S1**: The genetic load ($L$) and inbreeding load ($B$) for the gene and sites models, with varying values of the epistasis coefficient ($\epsilon$) when the rate of crossing over is low (one-tenth the standard value). Scaled selection coefficients followed a gamma distribution with $\bar{\gamma} = 100$ and shape parameter 0.3. 1000 selected sites were simulated with $Nu$ = 0.005 and $Nr$ = 0.001, where $N$ is the population size ($N$ = 1000), $u$ is the mutation rate per site and $r$ is the rate of crossing over between adjacent sites. Here $B$ was calculated by estimating the mean fitness of the population using 100 randomly sampled diploid individuals and is thus referred to as $B$ (sampled). The means and standard errors (SEs) for 1000 replicate simulations are shown.

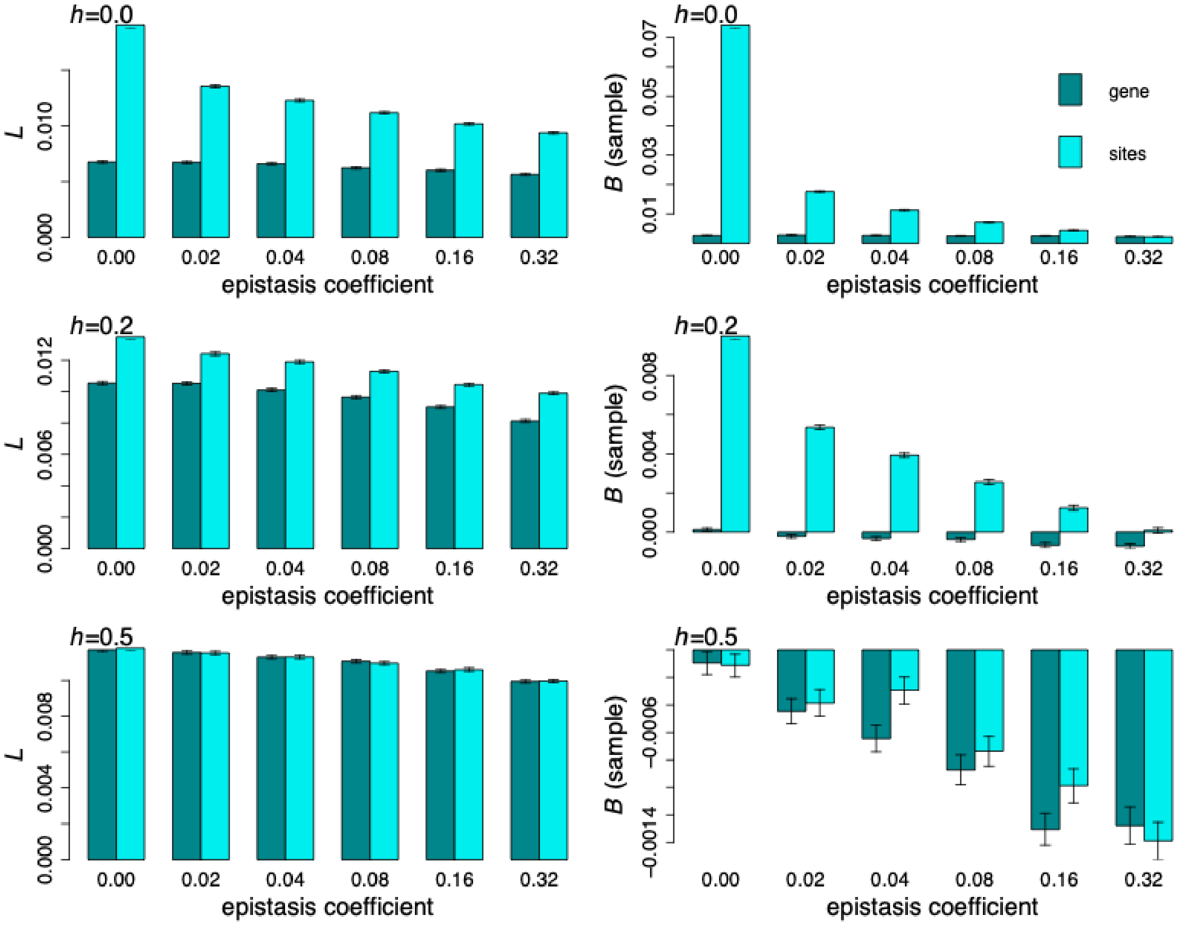

**Figure S2**: The variance in fitness ($Var$) and mean allele frequency ($q$) for the gene and sites models, with varying values of the epistasis coefficient ($\epsilon$) when the rate of crossing over is low. Simulations were performed as for Figure S1.

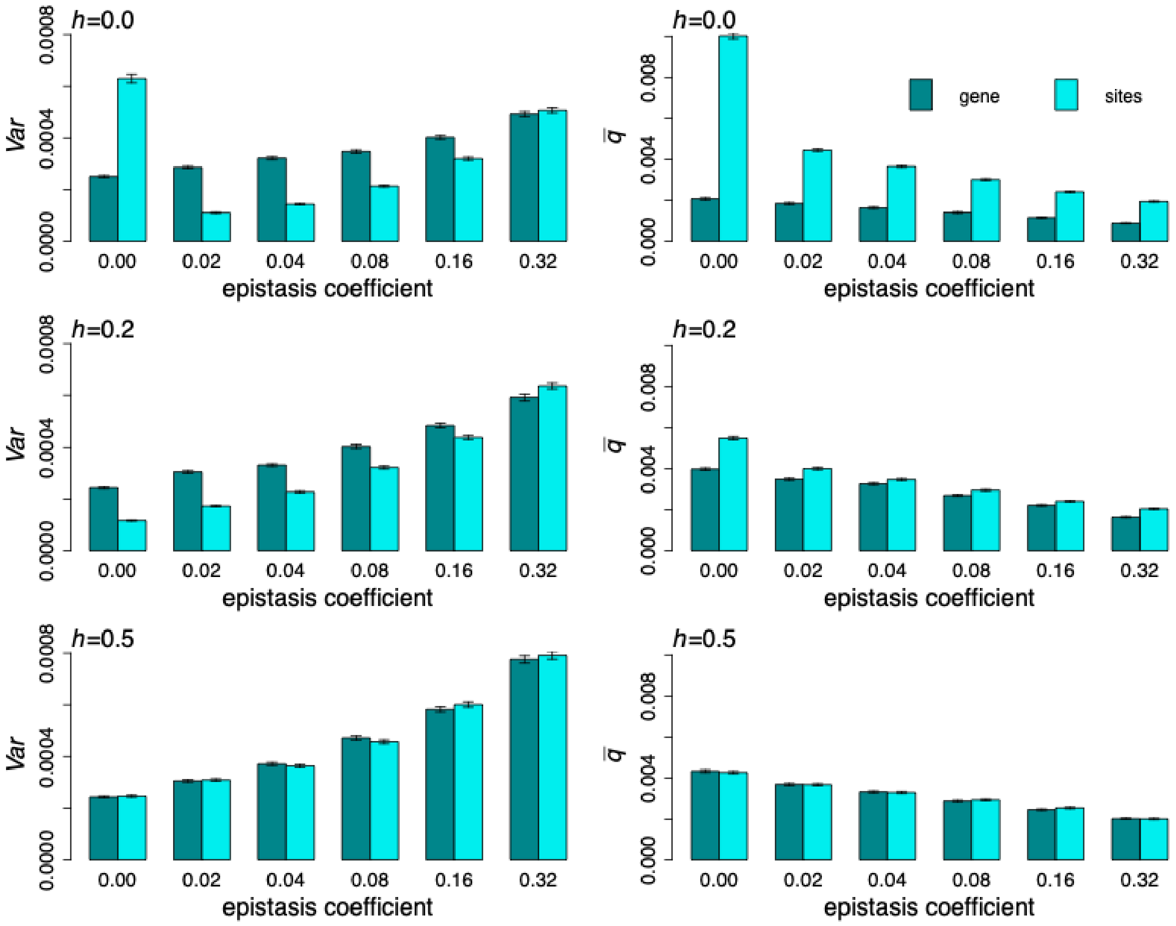
