## Supplementary File S2 for "A gene-based model of fitness and its implications for genetic variation: Genetic and inbreeding loads"

### FILE S2. POPULATION GENOMICS RESULTS ON *DROSOPHILA MELANOGASTER* RELEVANT TO THE INBREEDING LOAD

#### 1. Estimating the level of nonsynonymous diversity in *D. melanogaster* contributed by significantly non-neutral mutations

Campos et al. (2017) fitted a gamma distribution to data on the site frequency spectra of NS variants in a sample of genomes from a Rwandan population of *D. melanogaster,* together with divergence from *D. yakuba*. The method that was used (DFE-alpha of Eyre-Walker and Keightley 2009) assumes that mutations are semi-dominant ($h = 0.5$), for which the sites and gene models are equivalent in the absence of epistasis. The selection coefficient against a heterozygous mutation was thus assumed to be $t = ½s$, implying that their $\gamma = 4N_et$ is approximately equivalent to $4N_ehs$, given that selection against strongly selected mutations is predominantly against heterozygotes when mating is random. Their Figure S3 suggested an overall value of approximately 1250 for $2N_ehs$ for newly arising NS mutations, with a shape parameter of 0.3.

This estimate needs to be treated with caution, given that the method implicitly uses the sites model. If all or most mutations in coding sequences obey the gene model, then the fact that it predicts a substantially lower mean allele frequency than the sites model when $\gamma \geq 20$ (Table 3) implies that the strength of selection will be substantially overestimated by this method, and other similar population genomics approaches. The following analysis is therefore likely to overestimate the abundance of mutations that are under sufficiently strong selection that they behave quasi-deterministically, and hence will overestimate their contribution to the genetic and inbreeding loads.

A conservative threshold for mutations to behave deterministically is $2N_ehs \geq 2.5$ (Campos and Charlesworth 2019). Using numerical integration of the incomplete gamma function and the fact that $\Gamma(0.3) = 2.886$, this implies that the proportion of significantly selected new NS mutations is 0.91 (i.e., the proportion of NS mutations that are nearly neutral is 0.09) and yields a value of 0.0132 for the mean of $1/(2N_ehs)$ for such mutations.

From Campos et al. (2014), the mean synonymous site diversity, $\pi_S$, for autosomal loci in regions with normal recombination rates was 0.0141; with a mutation rate per basepair of approximately $4 \times 10^{-9}$, which is an approximate consensus value from the results in Table 1 of

Assaf et al. (2017; see also Wang et al. 2023). The corresponding value of $N_e$ is $8.81 \times 10^5$. There is probably some weak selection on these sites due to codon usage bias, and a value of $N_e = 10^6$ is a reasonable approximation to correct for this (Jackson et al. 2017). Using this value of $N_e$, together with the above estimate of the mean of $1/(2N_e hs)$ the mean of $1/(hs)$ for significantly selected NS sites is $2.64 \times 10^4$ (i.e., the harmonic mean of $hs$ at these sites is $3.78 \times 10^{-5}$). Under mutation-selection balance, the expected frequency of the mutant allele at a site is $q = u/(hs)$, and the expected mean diversity at significantly selected NS sites is then $\pi_{Ns} \approx 2\text{E}\{q\} = 2 \times 4 \times 10^{-9} \times 2.64 \times 10^4 = 2.11 \times 10^{-4}$.

A check on these calculations is as follows. The observed mean NS site diversity, $\pi_N$, was 0.00143. If the proportion of NS mutations that are neutral or nearly neutral is $p_n$, then $\pi_N \approx p_n \pi_S + (1 - p_n)\, \pi_{Ns}$. This gives $p_n = (\pi_N - \pi_{Ns})/(\pi_S - \pi_{Ns})$. The predicted value of $\pi_{Ns}$ is $1.06 \times 10^{-4}$, giving $p_n \approx (0.00143 - 2.11 \times 10^{-4})/(0.0141 - 2.11 \times 10^{-4}) = 0.088$, which agrees well with the population genomics estimate of 0.09 obtained above. A similar value is given in Figure 6 of Haerty and Ponting (2014).

#### 2. The mean number of slightly deleterious variants per *D. melanogaster* chromosome 2 and the mutation rate to non-lethal deleterious alleles

Using the above estimates, the mean frequency per nucleotide site of significantly deleterious NS variants is approximately one-half of $\pi_{Ns}$, i.e. $1.05 \times 10^{-4}$. Assuming a mean of 1500 exonic sites per gene, and a fraction of exonic sites that are NS equal to 0.7 (Campos and Charlesworth 2019), the mean number of significantly deleterious NS variants per gene in a random sample from the population is $1500 \times 0.7 \times 0.815 \times 10^{-4} = 0.110$. Given that chromosome 2 is 37% of the genome, and that there are approximately 14000 genes in the fly genome (Misra et al. 2002), the estimated mean number of significantly deleterious NS variants per 2nd chromosome is approximately 571.

This is much smaller than the estimate using $\pi_N$ itself, because the majority of NS variants segregating in the population are nearly neutral. If significantly selected non-coding variants are included, this number could be increased by a factor of two or even three, given that these contribute 3.8 times as much DNA as coding sequences and that approximately 50% of noncoding sites in *Drosophila* are under selective constraint (Halligan et al. 2006), although there is evidence for weaker selection against many non-coding sequence mutations than against NS mutations (Casillas et al. 2007; Haerty and Ponting 2014; Campos et al. 2017), reducing the fraction of segregating variants that are under significant selection.

The results of mutation accumulation experiments on *D. melanogaster* seem to require a small but significant contribution of mutations with homozygous selection coefficients that are much larger than those estimated from the population genomic data (Charlesworth 2015). Consensus values of the data from mutation accumulation experiments suggest a per 2$^{nd}$ chromosome mutation rate of 0.032 for large effect (but non-lethal) viability mutations. From Equation (A10) below, the corresponding value of $R$, the ratio of the mean fitness of non-lethal chromosome 2 homozygotes to that of heterozygotes, would be approximately 0.936 with a mutation rate of 0.032 and $h = 0.25$, not substantially different from 1.
