## Supplementary File S3 for "A gene-based model of fitness and its implications for genetic variation: Genetic and inbreeding loads"

### FILE S3. SOME FURTHER THEORETICAL RESULTS

#### 1. The validity of a cut-off of *$2N_ehs$* < 2.5 for nearly neutral mutations

A check on the use of the cut-off of $2N_ehs < 2.5$ as the criterion for mutations that are not under significant selection is provided by the simulation results in Table 4. With $\bar{\gamma}$ = 1000 and $h$ = 0.2, the gamma distribution with shape parameter 0.3 yields a proportion of mutations of 0.83 that exceed the cut-off. With a mutation rate of $0.5 \times 10^{-5}$ per site and 1000 sites in a gene, the mutation rate per gene adjusted for effectively neutral mutations is 0.005 x 0.83 = 0.00415. Using Equation (A10) with $h$ = 0.2, the $B$ value for a single gene of 0.00415 x 3 = 0.0124, compared with the simulation value of 0.0118 ± 0.0004 for the sites model. Similar calculations for $\bar{\gamma}$ = 100, 20 and 2 give the respective predicted $B$ values of 0.0091, 0.0055 and 0.0040; the corresponding simulation values for the sites model are 0.0098 ± 0.0001, 0.0071 ± 0.0001 and 0.00314 ± 0.0001. This simple approximation thus gives good fits to the simulation values for $B$, even for small mean values of $\bar{\gamma}$, accurately capturing the reduction in $B$ for partially recessive mutations when $N_es$ is reduced, as expected from previous theory (e.g., Charlesworth 2018).

This calculation can be made even simpler by noting that the cumulative gamma distribution using the scaled variable $y = \alpha\gamma/\bar{\gamma}$ for $y << 1$ between 0 and $y_0$ is approximated by:

$$\Phi(y_0) \approx \Gamma(\alpha)^{-1} \int_0^{y_o} y^{\alpha-1}\,\mathrm{d}y = \Gamma(\alpha)^{-1}\alpha^{-1}y_0^{\alpha} \qquad \text{(S3.1)}$$

If $\alpha << 1$, $\Gamma(\alpha) \approx \alpha^{-1}$, so that:

$$\Phi(y_0) \approx y_0^{\alpha} \qquad \text{(S3.2)}$$

With $\alpha$ = 0.3, this approximation is quite accurate even for $y_0$ as large as 0.25, so that $1 - y_0^{0.3}$ can be used for most practical purposes to determine the proportion of sites that are under significant selection, with the appropriate choice of $y_0$.

**2. Why is the inbreeding load reduced when $N_e s$ is small?**

For the results presented in Tables 3 and 4, the reduced level of variability in small populations must be involved in the reduction in $B$ with smaller $N_e s$, since we are comparing different strengths of selection while keeping $N_e$ and the mutation rate constant. In Table 4, as expected from the effects of reduced $s$, $\bar{q}$ changes from 0.0092 ± 0.0028 with $\bar{\gamma}$ = 1000 to 0.0383 ± 0.0053 with $\bar{\gamma}$ = 2, the opposite of what is expected with purging; the ratio of $\bar{q}$ values (equal to 4.16) is, however, much less than the 50-fold difference in $\bar{\gamma}$. A similar pattern of decrease in $\bar{q}$ with increasing $\bar{\gamma}$ is seen with $h$ = 0.5. The explanation for the lack of proportionality between the two variables is explored in section 3 below.

In contrast to $B$, the genetic load and variance in fitness are little affected by the effects of drift when selection is weak. This is illustrated by the results on the genetic loads under the sites model for different $\bar{\gamma}$ values and $h$ = 0.2 in Table 4; the mean loads for $\bar{\gamma}$ = 1000 and $\bar{\gamma}$ = 2 are 0.0101 ± 0.001 and 0.0095 ± 0.001, respectively, and the approximate deterministic value of $2000u$ (Haldane 1937) is 0.01. Even with $\bar{\gamma}$ = 2, the load is close to the deterministic value, in contrast to the sizeable reduction in $B$. The variances in fitness under the sites model for $h$ = 0.2 and 0.5 in Table 4 are also well approximated by the deterministic value of $4000uh\bar{s}$.

**3. The relation between mean $q$ and $\gamma$**

The fact that the mean frequencies of deleterious alleles are not inversely proportional to the strength of selection when $\gamma$ is relatively small can be understood by examining the case of $h$ = 0.5 and a fixed value of $\gamma$ (see Table 3). In the absence of LD, the predicted value of $\bar{q}$ with our scaled mutation rate of $4Nu$ = 0.02 with $\gamma$ = 2 should be closely approximated by the Li-Bulmer equation, which predicts the proportion of sites fixed for the favorable allele in a diallelic mutation-selection model (McVean and Charlesworth 1999). With no mutational bias, this gives:

$$\bar{q} \approx \frac{\exp(-\gamma)}{1+\exp(-\gamma)} \qquad \text{(S3.3)}$$

With $\gamma$ = 2, $\bar{q} \approx$ 0.12, compared with the simulation value of 0.0236 ± 0.0039 for the sites model. For $\gamma$ >> 1, $\bar{q}$ is approximated by the deterministic value $u/(hs) = 2u/s$, which is equal to 0.0013 with $\gamma$ = 20, compared with 0.00079 ± 0.00014 for the sites model. The observed ratio of

$\bar{q}$ values is 30, compared with a predicted range of 0.12/0.0013 = 92. This suggests that selection is more effective in the multi-site context than for a single site in isolation when selection is relatively weak. At first sight, this is puzzling, because Table 2 of Johri and Charlesworth (2025) shows no evidence for significant LD with $h$ = ½.

The probable explanation is as follows, which is based on the two-sites model (Table 1). Most of the contributions to $\bar{q}$ with these parameters come from fixed rather than segregating sites (McVean and Charlesworth 1999). Any effects of LD on $\bar{q}$ come from transitions between fixed pairs of sites where two variants are segregating simultaneously. Transitions of the type + – to – – and + + (or vice versa) are likely to only temporarily involve two simultaneous segregating sites, and can be ignored as far as the effects of LD on the outcome of selection are concerned, although they are the most common type of transition and source of variability. Transitions of the type + + to – – must involve a first mutation of the type + + to + – , followed by a + – to – – mutation; in the absence of recombination, this creates a two-site polymorphism during the transition of the type discussed in the section *Effects of weighting towards low frequencies on LD statistics* in the companion paper Johri and Charlesworth (2025). This has $D > 0$ when the sign of $D$ is defined by selective status. This enhances the efficacy of selection, and hence reduces the chance of this transition. Similarly, transitions of the type – – to + + have $D > 0$, and their probability of success is thereby increased over the single-site expectation. Thus, despite the lack of overall positive LD, the tendency for selection to favor + versus – alleles is enhanced by linkage, thereby reducing $\bar{q}$ below single-locus expectation. This effect is likely to be largest when selection is relatively weak, when transitions of these two types are relatively common, explaining why the ratio of $\bar{q}$ values for weak versus strong selection is less than might be expected.
